# An atlas of protein homo-oligomerization across domains of life

**DOI:** 10.1101/2023.06.09.544317

**Authors:** Hugo Schweke, Tal Levin, Martin Pacesa, Casper A. Goverde, Prasun Kumar, Yoan Duhoo, Lars J. Dornfeld, Benjamin Dubreuil, Sandrine Georgeon, Sergey Ovchinnikov, Derek N. Woolfson, Bruno E. Correia, Sucharita Dey, Emmanuel D. Levy

**Author notes:** Emails (ORCID).

## Abstract

Protein structures are essential to understand cellular processes in molecular detail. While advances in AI revealed the tertiary structure of proteins at scale, their quaternary structure remains mostly unknown. Here, we describe a scalable strategy based on AlphaFold2 to predict homo-oligomeric assemblies across four proteomes spanning the tree of life. We find that 50% of archaeal, 45% of bacterial, and 20% of eukaryotic proteomes form homomers. Our predictions accurately capture protein homo-oligomerization, recapitulate megadalton complexes, and unveil hundreds of novel homo-oligomer types. Analyzing these datasets reveals coiled-coil regions as major enablers of quaternary structure evolution in Eukaryotes. Integrating these structures with omics data shows that a majority of known protein complexes are symmetric. Finally, these datasets provide a structural context for interpreting disease mutations, which we find enriched at interfaces. Our strategy is applicable to any organism and provides a comprehensive view of homo-oligomerization in proteomes, protein networks, and disease.

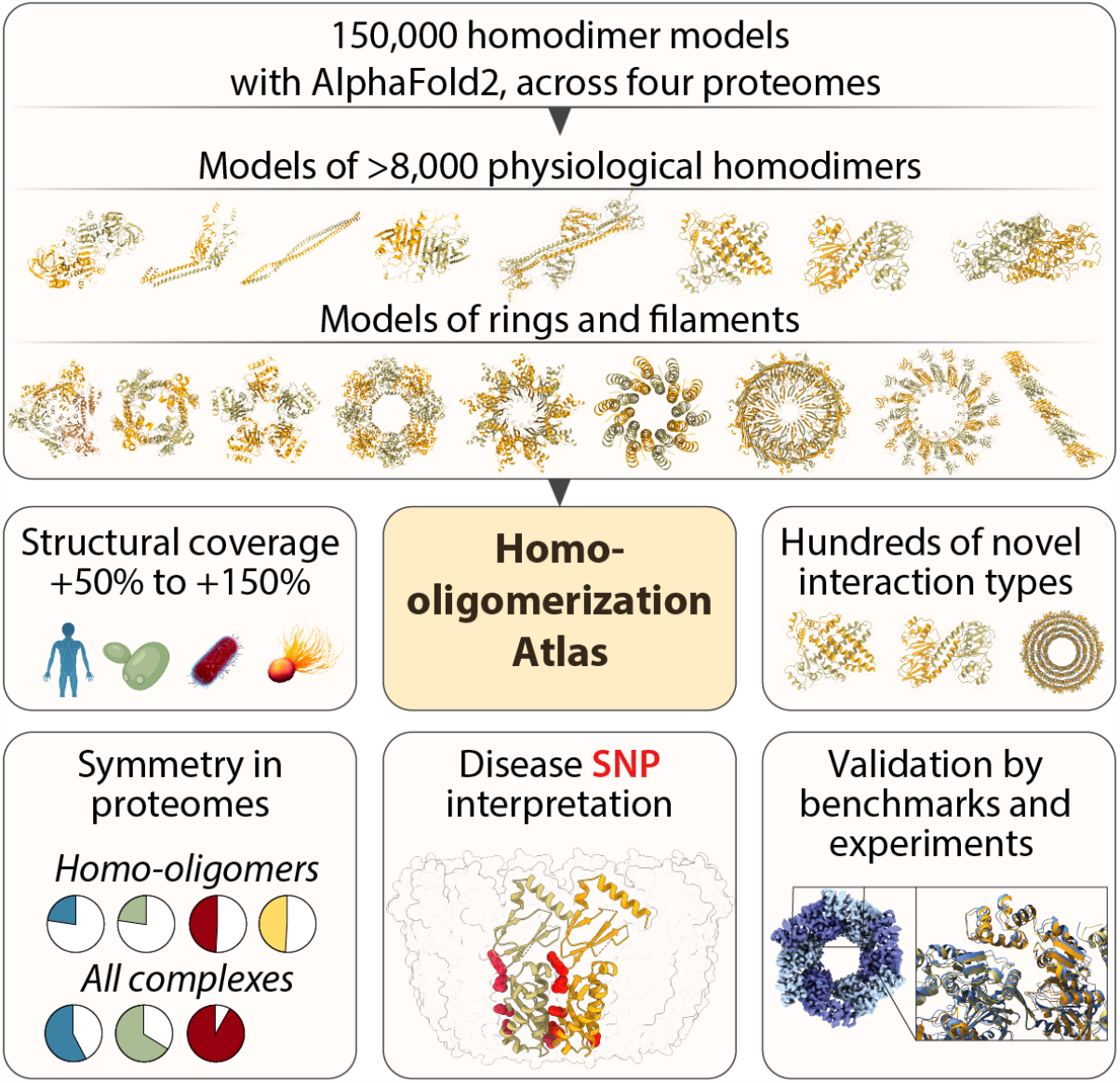

## Introduction

The organization of proteins into complexes and biomolecular networks underlies cellular processes and functions. At the most fundamental level, protein assembly occurs by homo-oligomerization, whereby identical copies of a protein interact symmetrically to form higher-order structures ^1,2^. These so-called homomers possess unique structural and functional properties ^1^ (**Fig, 1A**). They enable forming repetitive structural elements, as in the cytoskeleton, they can create unique shapes like rings, barrels, or cages ^3^. More broadly, the repetition of protein chains in homomers provides multivalence, a parameter critical to the formation of biomolecular condensates ^4^. Functionally, the conformation of their subunits can be coupled and mediate allosteric transitions, and their formation can be modulated by environmental cues, such as pH or post-translational modifications ^5^. As such, comprehensive knowledge of homomer structures provides a foundational layer of information to analyze and interpret protein structure and function. In particular, it would allow modeling and predicting the underlying molecular basis of a wide variety of human diseases and associated mutations that occur at, or close to interfaces. For example, Aquaporin forms ring-like channels allowing water to flow through membranes, and mutations impairing the ring assembly are associated with nephrogenic diabetes insipidus disease ^6^. Beyond their functional importance, homomers are shaping the evolution of protein complexes and networks: they represent the ancestral state of myriad key macromolecular complexes such as histones, proteasomes, or chaperones, which diversified through gene-duplication ^7^.

**Figure 1.**
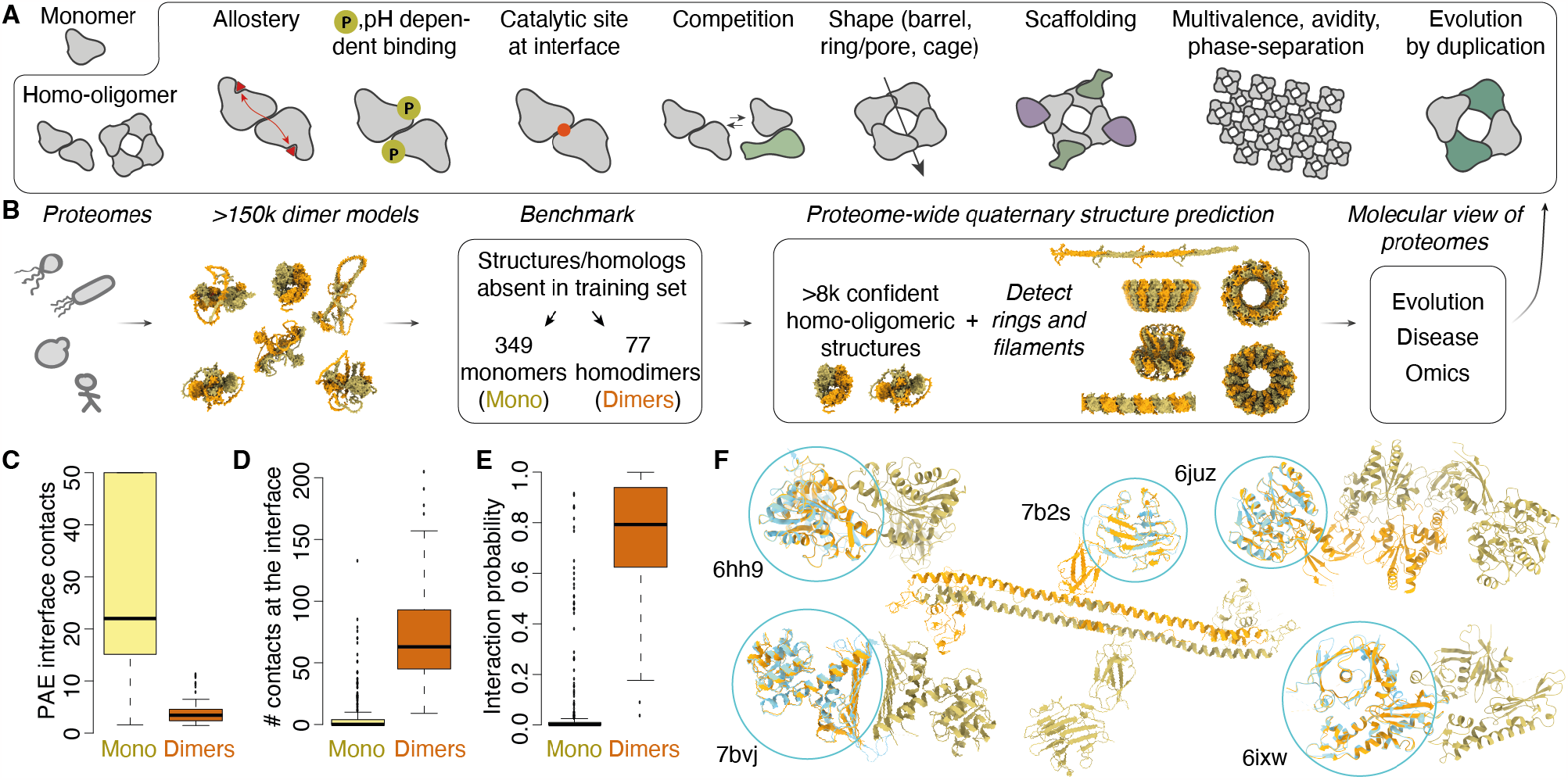
AlphaFold2 predicts the structure of homodimers with high accuracy. **A**. Homo-oligomeric proteins possess unique functional, shape, and evolutionary properties. **B**. Overview of the data flow in this work. Dimer structures were predicted for proteins across four species’ proteomes, yielding 156,065 models. The models were subsequently scored based on a benchmark, yielding over 8,000 high-confidence dimer structures. The dimer structures were used to predict higher-order biologically relevant macromolecular assemblies into rings and filaments, yielding proteome-wide homo-oligomerization information. **C**. The position alignment error (PAE) of contacting residues discriminates dimers from monomers. **D**. The number of contacts between subunits also discriminates dimers from monomers. **E**. The features from panels B and C were combined by logistic regression. Benchmark data are detailed in **Data S1. F**. Examples of discrepancies where the experimental structure is a monomer (blue and circled, PDB code indicated) while our predictions suggest a dimer (orange/green).

The central role of homomers in biology motivates their comprehensive characterization. Recent advances in machine learning have revolutionized the accuracy with which protein tertiary structure is predicted ^8,9^. These advances have been scaled up, making the structure of monomeric proteins available across entire proteomes ^10,11^. Machine learning approaches can also be employed to predict the structure of protein complexes ^12,13^ and served to predict heteromeric complexes in yeast ^14^ and human ^15^. However, two major challenges make it difficult to predict homo-oligomers systematically, on a proteome-wide scale. While AlphaFold2 has been the method of choice, it requires knowledge of the number of protein copies present in a complex, and this number is typically unknown. Additionally, computation and memory requirements scale exponentially with the number of copies modeled by AlphaFold2, making it difficult to predict large complexes at scale.

Here we addressed these challenges to predict the structure of homo-oligomers on a proteome scale. We systematically generated structures for putative homodimers and analyzed them to identify those with physiological relevance. The latter were subsequently processed independently of AlphaFold2 to predict higher-order structures, including rings and filaments. We computed homo-oligomeric structures for four species: *P. furiosus, E. coli, S. cerevisiae*, and *H. sapiens*. The resulting datasets comprise 872, 2181, 1196, and 3946 homo-oligomers, covering 20-40% of the analyzed proteomes. This emphasizes that a considerable fraction of the proteome undergoes homo-oligomerization, highlighting once more the importance of this phenomenon for understanding protein structure, function, and evolution. A number of these models recapitulate large structures including a hexameric ring that we validated experimentally by cryo-EM, or a megadalton macrophage pore-forming complex, which consists of a ring with 16 protein copies ^16^. These datasets add a quaternary structure dimension to proteomes and will bolster our molecular understanding of their function and evolution (**Fig. 1B**). We illustrate such new biological insights in three analyses showing that: (i) coiled-coil regions are major enablers of quaternary structure evolution in eukaryotes; (ii) interaction interfaces in homo-oligomers across the human proteome are 70% more likely to contain disease mutations than protein surfaces; and (iii) strikingly large fractions of homo and hetero-oligomeric protein complexes in prokaryotes and eukaryotes are symmetric.

## Results

### Predicting homodimers with AlphaFold2

We first assessed the accuracy of AlphaFold2 at identifying homodimers and correctly predicting their structure. We used the initial AlphaFold2 weights rather than the “multimer” weights because the gain in accuracy for homo-oligomers appeared limited ^12^. Additionally, those weights were trained on single chains, thus avoiding overfitting when predicting multi-chain interactions in homo-oligomers. We compiled a non-redundant dataset from the PDB ^17^ consisting of 349 monomers and 77 homodimers with a structure deposited after June 2018 (**Fig. 1B**, Methods). Predictions of multiple metrics showed an excellent agreement with the X-ray crystallography-derived dataset of monomers and dimers (**Fig. S1, Data S1**). Two metrics were particularly informative in discriminating physiologically relevant homodimers from monomers: the first is the average predicted alignment error (PAE) of amino acids in contacts (**Fig. 1C**), and the second is the number of contacts between amino acids (**Fig. 1D**). We combined both metrics in a logistic regression model, which predicts the probability of an AlphaFold2 dimer to be physiologically relevant (**Fig. 1E**). This simple two-parameter model accurately captured the oligomeric state of experimental crystal structures, with an area under the receiver operator curve (AUC) of 0.978. Manual inspection of the predictions revealed several cases where the AlphaFold2 model appeared physiologically relevant (**Fig. 1F**). In the case of a two-domain esterase (PDB code 6HH9 ^18^), the predicted dimer was, in fact, observed experimentally and existed in the crystal lattice, hinting at its possible existence in solution. In several instances (e.g., PDB codes 7B2S, 6JUZ ^19^), the structure solved by X-ray crystallography was truncated and did not include the dimerization domain, which explained the apparent inconsistency. In another example (PDB code 7BVJ ^20^), the primary reference provided evidence for the formation of a homodimer. In the last example, an actin-like protein appeared in the PDB as a monomer due to point mutations, while the physiological interaction interface driving filament assembly was detected by AlphaFold2 despite these mutations.

Overall, this analysis shows that AlphaFold2 accurately predicts the structure of homodimers and that we can efficiently discriminate between physiological homodimers and artifactual complexes. These results motivate the generalization of its use to discover homo-oligomers across proteomes.

### Proteome-wide discovery of homodimers

Protein structure inference is computationally expensive and hardly applicable to large complexes on a proteome-wide scale. To address this limitation, we adopted a hierarchical approach where we initially predicted homodimers, and subsequently analyzed whether they form larger structures based on the dimer’s internal symmetry. We generated a total of 156,065 homodimer models altogether covering 99.8%, 98.2%, 94.7%, and 89.7% of reference proteomes ^22^ for *P. furiosus, E. coli, S. cerevisiae, and H. sapiens* respectively.

We analyzed the inter-subunit contacts of these models, their consistency across the five AlphaFold2 networks, as well as their predicted aligned error (PAE) statistics. We then used these metrics (**Fig. 1, Fig. S1**) to calculate confidence probabilities based on the benchmark set. The scoring process (Methods) yielded 872, 2181, 1196, and 3946 homodimers that covered 43%, 44%, 21%, and 21% of the four proteomes, respectively (**Fig. 2A**, detailed information about reference sets is provided in **Data S2**). A significant fraction of these predictions closely matched sequences with an experimentally solved structure. As we employed a version of AlphaFold2 trained exclusively on single chains, we evaluated whether these models recapitulated known homo-oligomeric structures. This comparison revealed an excellent agreement, with 95.3%, 97.7%, 98.9%, and 98.7% of models recapitulating the known interaction interface among *P. furiosus, E. coli, S. cerevisiae*, and *H. sapiens* respectively (**Fig. 2C, Fig. S2**). We did not find structural homologs for 15-20% of the models, which thereby represent hundreds of potentially new quaternary structure types (**Fig. 2B**). While the pace of new protein fold discovery is relatively slow, likely due to the extensive coverage of existing structures, the number of quaternary structure types we discovered indicates that they cover a vast structural landscape, much larger than that of tertiary structures (**Fig. 2B**).

**Figure 2.**
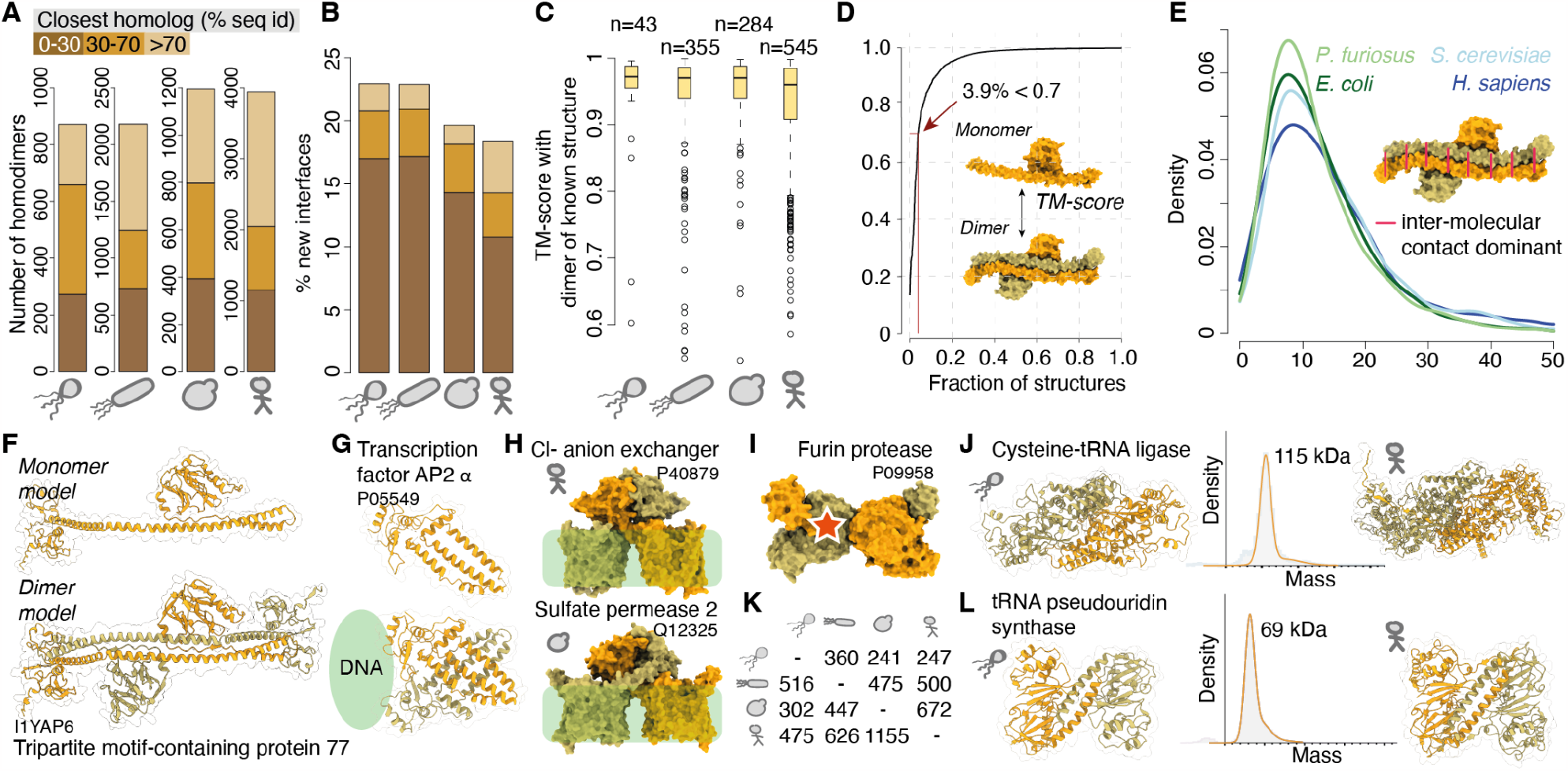
Proteome-wide discovery of homodimers. **A**. Numbers of high-scoring dimers in each species’ dataset and their similarity to known structures. **B**. Fraction of dimer models with no structural homolog detected in the PDB. **C**. Structural similarity between dimer models and the matching experimental structure when available. Outliers are detailed in **Fig. S2. D**. Structural similarity of protein chains across models inferred separately as monomers or dimers. **E**. Dimers often contain significant intermolecular contacts that stabilize the structure. **F**. The monomer (left) and dimer (right) models of the Tripartite-motif-containing protein 77. This dimer’s extensive intermolecular contacts highlight the necessity to consider oligomerization information for interpreting the new wealth of structural data. **G**. The transcription factor AP-2 α functions as a dimer ^21^ but its structure is unresolved. Our dimer model represents a new quaternary structure type (i.e., not found in the PDB) and is compatible with biochemical data ^21^. **H**. Membrane transporters that adopt a quaternary structure type shared between human and yeast, and absent from the training set. **I**. The dimer model of the furin protease shows a pro-form that trans-inactivates. The red star indicates the catalytic site obstructed by binding of the partner. **J**. A cysteine tRNA ligase shows a novel dimer interaction geometry conserved between archaea and eukarya and absent from the PDB. The purified protein from archaea (Q8U227) showed a scattering peak at 115 kDa, confirming its existence as a dimer in solution (this panel is detailed in **Fig. S3**). **K**. Number of dimers from each species (line) sharing structural homology with dimers from the other species (columns). **L**. A tRNA pseudouridine synthase shows a novel and similar dimer interaction geometry in archaea and eukarya. The purified protein from archaea (Q8U2C1) showed a scattering peak at 69 kDa, compatible with a dimer state in solution (this panel is detailed in **Fig. S3**).

Next, we focused on obligate homo-oligomers, which are expected to be unstable as monomers ^23^. We reasoned that predicting monomers instead of dimers could reveal such complexes, as we assumed that their structure would change between both states due to their instability as monomers. We generated monomer models for the human proteome and compared the resulting structures to those of individual chains in dimers. Surprisingly, we observed a high similarity in structure between chains in either state, with less than 4% of homodimer chains showing a structure highly different (TM-score < 0.7) from that of the monomer (**Fig. 2D**). Such a degree of similarity was unexpected because a large fraction of our dataset corresponds to structures with dozens of residues stabilized by intermolecular contacts (**Fig. 2E**). This observation shows that AlphaFold2 identifies native-like structures of protein chains forming obligate complexes, even in the absence of their partner.

The substantial number of proteins displaying extensive intermolecular interactions underscores the need to represent these models in their quaternary dimension. This is particularly crucial for effectively utilizing the recent expansion of structural descriptions within the proteome and ultimately for understanding their biological significance. For example, the Tripartite motif (TRIM)-containing protein 77 (**Fig. 2F**) is stabilized by considerable inter-subunit contacts, yet the monomer chain exhibits a similar structure as in the dimer. Moreover, dimer information is key to visualizing the multivalent and spatial organization of the RING and SPRY domains in this protein, information that is absent from the monomeric structure. In a different example, the transcription factor AP-2-α (**Fig. 2G**) exhibits extensive intermolecular contacts. This family of helix-span-helix transcription factors has no experimentally determined structure, and accordingly, this quaternary structure type is novel with no homologs detected across the PDB. Moreover, this dimer matches existing biochemical and mutational data ^21^, providing further validation of its accuracy. We also identified novel quaternary structure types among membrane proteins. For example, the chloride/bicarbonate anion exchanger S26A3 shows an interface geometry absent from the training dataset, but is substantiated by the recent characterization of a homolog ^24^. Interestingly, the proteome-wide nature of our predictions enables comparing these structures across organisms and revealed a homologous sulfate transporter in *S. cerevisiae* (**Fig. 2H**).

Our models also pinpoint new potential regulatory features difficult to grasp experimentally. For example, the furin protease is a key enzyme that processes cellular precursor proteins and viral factors essential for the function of HIV, influenza, and SARS-COV-2. This protease must remain inactive in its intra-cellular form to avoid mis-cleavage events, but the structure of its proform is unknown. Our models suggest that furin and other family members, including in archaea, trans-inactivate as dimers whereby each chain binds and obstructs the catalytic site of its partner (**Fig. 2I**).

In a different example, we identified a cysteine tRNA ligase as exhibiting a new quaternary structure type conserved across archaea and eukarya, and the purified protein from archaea exhibited the expected dimer molecular weight in solution (**Fig. 2J, Fig. S3A**). The comprehensive nature of our datasets renders them suitable to analyze homo-oligomerization conservation and evolution. We found that 247 and 500 dimer structures from archaea and bacteria respectively shared structural homology with dimers from the human proteome. Conversely, 475 and 626 dimers in the human proteome shared structural homology to a dimer from archaea or bacteria respectively (**Fig. 2K**). These data can also serve to identify cases of divergence and will provide a basis for comprehensive analyses of interface and oligomeric state evolution. One notable example is the tRNA pseudo-uridine synthase. This protein adopts a novel quaternary structure type observed in the proteomes of archaea, yeast and human, and the purified protein from archaea exhibited the expected dimer state in solution (**Fig. 2L**). Interestingly, in E. coli, the interaction interface occurs at a similar surface site but is mediated by extended loop regions absent in other species (**Fig. S3B**).

Taken together, these analyses show that these novel structure models are reliable, they increase the structural coverage of homo-oligomer information by ∼50% in human to >100% in *P. furiosus*, they contain hundreds of new quaternary structure types, and therefore provide a rich resource for functional and evolutionary analyses.

### Proteome-wide discovery of ring and filament-forming homo-oligomers

A majority of homo-oligomers form homotypic or “head-to-head” interfaces, resulting in dimers with C2 symmetry. A different type of assembly involves heterotypic or “head-to-tail” interactions, which create ring structures. These rings are difficult to predict due to the uncertainty in the number of subunits, and due to their large size. However, we reasoned that the symmetry information contained within a dimer could suffice to reconstruct ring-like and filament-forming complexes. This concept is illustrated with the synporter SatP, where the predicted dimer model interacts head-to-tail. The rotation information contained within the dimer is best compatible with C6 symmetry, which we identified through an analytical method^25^ (**Fig. 3A**). This strategy yields a model of SatP closely matching the experimental structure (TM-score = 0.99). Comparing the symmetries derived with this approach to their matching experimental structures also reveals an excellent agreement, with 95% (160/168) of cyclic symmetries being inferred correctly (**Fig. 3B**). This strategy thus tackles both limits of Alphafold2, first by inferring the number of subunits given the symmetry of a dimer, and second by decreasing the resources required for predicting these complexes. Indeed, we compared time and memory requirements for predictions with AlphaFold2 “Multimer” ^12^. Complexes with 4 to 10 subunits required 5 to 50-fold more time (**Fig. 3C, Fig. S4**) and 1.3 to 6.5-fold more GPU memory than for their respective dimer predictions.

**Figure 3.**
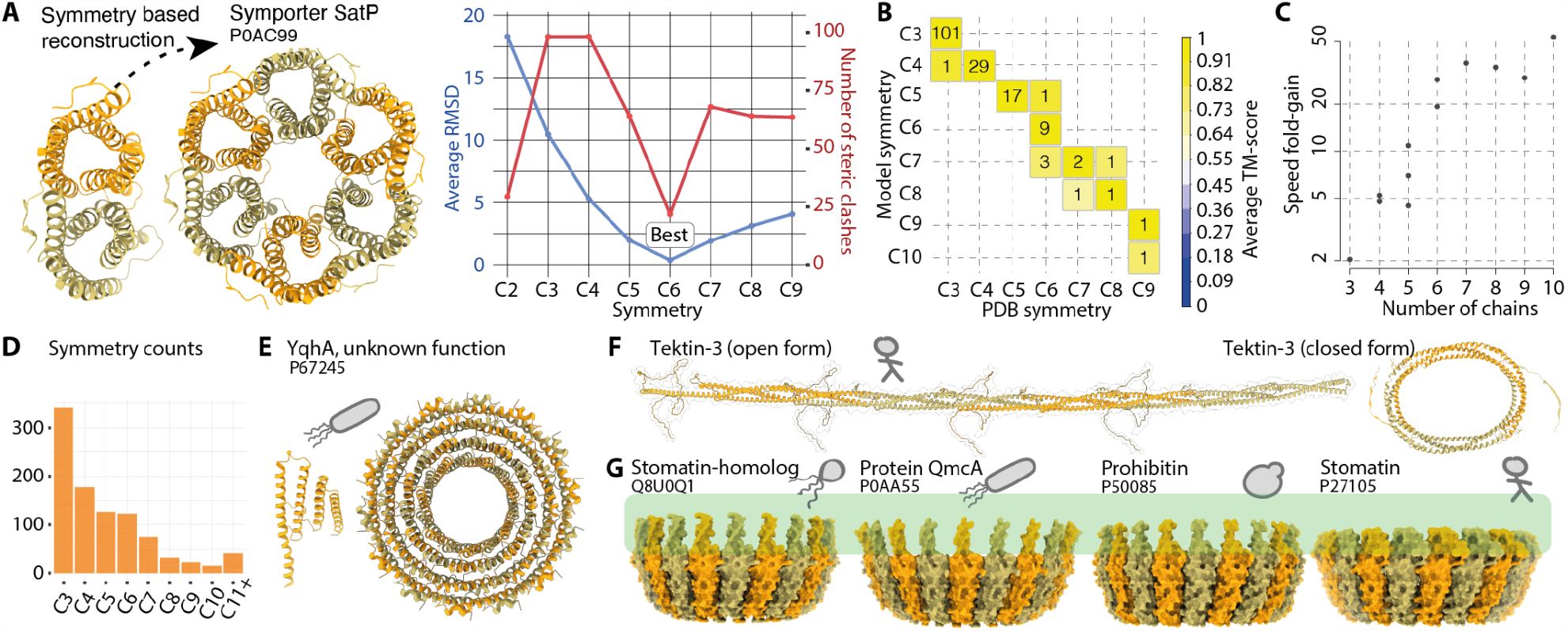
Proteome-wide discovery of ring complexes and filaments. **A**. The symmetry information contained in a dimer model is used to find the best-compatible ring symmetry. **B**. The structure of cyclic complexes so obtained was compared against known experimental structures, showing that 95% (160/168) of cyclic symmetries are inferred correctly. The average TM-score is above 0.93 for all bins on the diagonal and is above 0.7 for off-diagonal predictions, except for the C8-C7 bin (0.69). **C**. Speed fold-change observed in the prediction of complexes with containing three to ten chains. **D**. Numbers and types of ring symmetries reconstructed across the four proteomes. **E**. A monomer of Yqha is shown next to its ring structure, which represents a novel quaternary structure type. **F**. Tektin-3 is predicted as forming a filamentous structure identical to that recently proposed ^30^ and a novel, closed form is also predicted. **G**. Stomatin and prohibitin proteins are membrane-associated. They assemble as rings through an interface geometry conserved across kingdoms.

One drawback of the symmetry-based reconstruction of ring complexes is that loops could be intertwined and flexible regions sometimes clashed extensively with the ring structure. To overcome this problem, we developed a new AlphaFold2 protocol that makes use of the backbone of the complete symmetry-generated structure to produce final models. We supplied the structural information to AlphaFold2 as a template of which the sidechain information had been masked, limiting a too heavy bias towards the generated structure. However, we found that many recycles were needed and sometimes not sufficient for large structures. We hypothesize that this was due to the “black hole” initialization of the structure module, where atomic coordinates are all initialized at zero. Hence, we implemented what we call a “big bang” initialization, where instead of initializing the coordinates at zero, we initialized them to the input structure, resulting in faster and consistent convergence to the final model (Methods). This protocol allowed us to reconstruct the ring complexes with up to 6,500 residues in total while resolving the clashes introduced by the symmetry-based model generation.

This strategy enabled us to reconstruct hundreds of ring complexes (**Fig. 3D, Fig. S5**), many of which represent entirely novel structures. One example is Yqha, a protein of unknown function from *E. coli*. The monomer structure of this protein consists of four helices interacting laterally, which appears highly unstable due to the absence of a protein core. By contrast, our model shows how the four helices pack with additional copies to form a ring structure containing 14 subunits (**Fig. 3E**). In a different example, we noticed an unusual structure of intertwined alpha-helices for caveolin-2 (**Fig. 4A**) closely resembling that of caveolin-1 solved by cryo-EM ^26^ and absent from AlphaFold2 training set. This example motivated us to evaluate the accuracy of the models against human homo-oligomers specifically solved by cryo-EM. The models matched these structures closely, as illustrated for the transmembrane protein 45A (TM-score=0.95, **Fig. 4B**), or the megadalton complex of macrophage expressed gene 1 (TM-score=0.98, **Fig. 4B**). Overall, out of 20 complexes so compared (**Data S3**), the median TM-score was 0.92, reflecting an excellent agreement (**Fig. 4C, Fig. S6**). Most of the mismatches between models and experimental structures were caused by differences in the cyclic symmetry we inferred (e.g., C12 instead of C11 in the case of the human calcium homeostasis modulator 5, (PDB code 7d60 ^27^). Such symmetry changes involve minute structural differences in the dimer interaction geometry and proteins can frequently adopt multiple states. This is also observed among viruses adopting a quasi-symmetry ^28^. Indeed, focusing the benchmark on the dimer interaction geometry, our prediction accuracy for the same structures increases significantly (**Fig. 4C**, olive).

**Figure 4.**
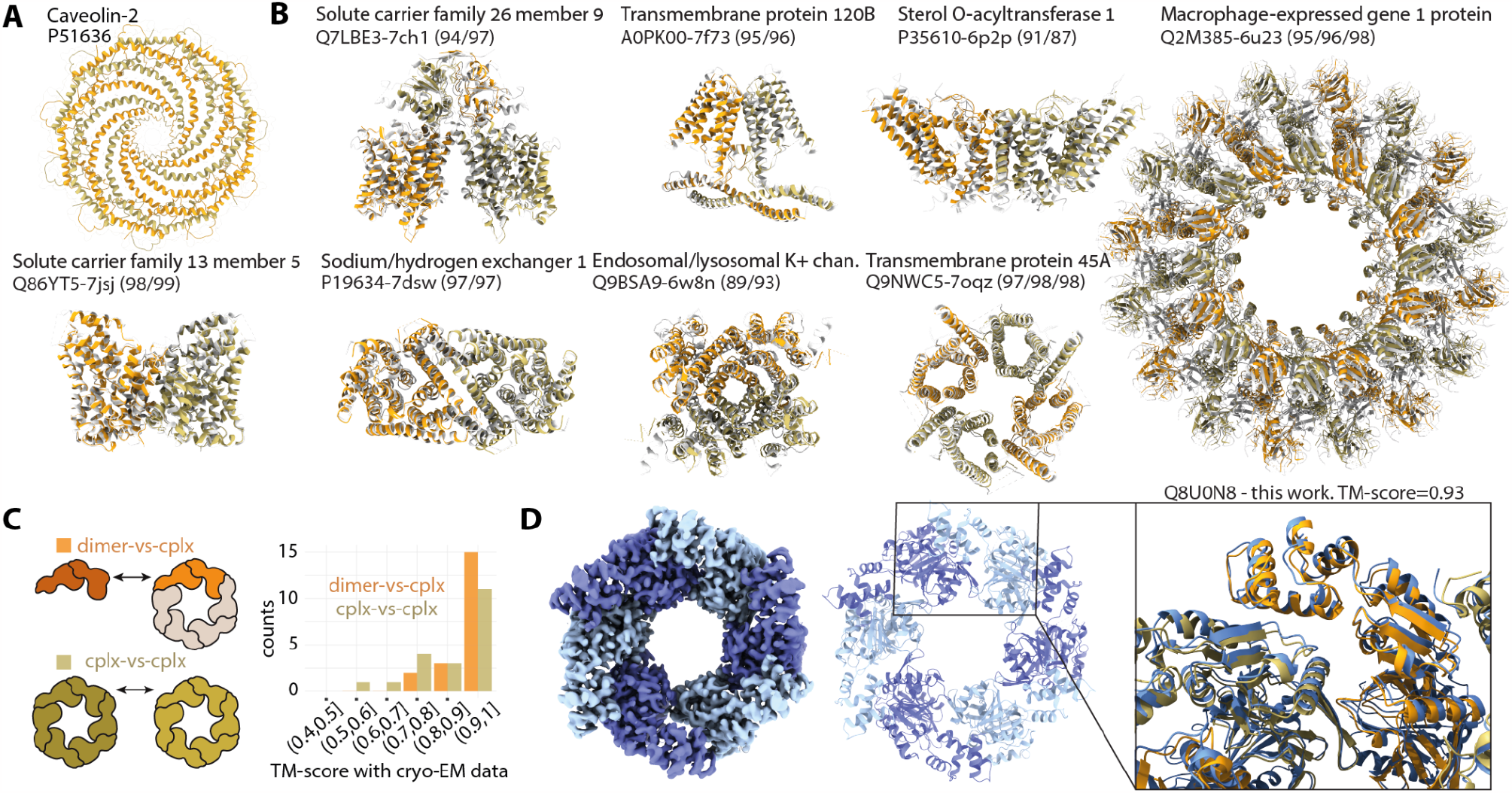
Assessing the accuracy of predictions based on recent cryoEM structures. **A**. The model of caveolin-2 is nearly identical to a recently published structure of caveolin-1, absent from the training set ^26^. **B**. Models of cyclic complexes (orange-green) superposed to cryo-EM structures (white) released after May 2018 and with no close homolog in the training set. The name and Uniprot ID of the proteins are given along with the PDB code of the matching structure. Numbers in parenthesis indicate the TM-score*100 for the chain/dimer/complex superposition. **C**. Histogram of TM-score between the models and 20 such EM structures. **D**. Using cryoEM, we solved the hexameric structure of a protein of unknown function from *P. furiosus*. Our model captured the quaternary structure of this complex with high accuracy. Left shows cryoEM density, middle the corresponding atomic model, right the overlay between the experimental structure (blue) and the predicted model (orange).

To validate our predictions, we selected several ring-forming proteins for structural determination. One of them, which was predicted to form a hexamer, could be expressed and was amenable to analysis by cryo-EM. This protein (Q8U0N8) is uncharacterized and does not have a PFAM domain assignment ^29^. We solved its structure to a resolution of 2.8 Å, which revealed a close agreement with our prediction. The global TM-score between our model and the determined structure was 0.93 (**Fig. 4D**). Interestingly, the protein contains an N-terminal domain that exhibits varying degrees of flexibility, as indicated by the reduced local resolution (**Fig. S7**). Not considering this domain, our model showed higher agreement with the experimental data (TM-score=0.96, Cα-RMSD = 2.4 Å).

Beyond ring complexes, we also identified 179 models expected to form filamentous assemblies (**Figure S3, Data S2**). One such example is Tektin-3, a component of dynein-decorated doublet microtubules. Here, the filament structure is identical to that recently proposed ^30^. Interestingly, one of the five models of Tektin-3 converged towards a different conformation corresponding to a homodimer. We speculate that such a closed structure could be adopted after synthesis, to facilitate delivery to doublet microtubules. Lastly, we also identified one novel quaternary structure type conserved in all kingdoms. The proteins forming these structures were annotated with an “ambiguous” or “Trans” symmetry (**Fig. S3**) because they contain a flexible coil conflicting with the symmetry search procedure. However, upon truncating that region, the procedure predicts ring-shaped assemblies containing about 20 chains and compatible with negative-stain images of yeast prohibitin ^31^. In human, Stomatin and homologous proteins such as Podocin associate with lipid rafts and diverse ion channels to regulate their activities ^32,33^. The molecular basis of these interactions remains unknown despite their association with numerous diseases ^34^, and the ring-shaped assembly characterized here provides a structure with which they can be interpreted, as we will show in the next section.

### Evolutionary and structural insights from proteome-wide homo-oligomerization

The proteome-wide characterization of quaternary structures paves the way to a molecular description of proteomes, both in health and disease. We employed the newly characterized datasets to investigate three general molecular properties of proteomes.

First, we clustered the dimer models by structural similarity to identify the most frequent type of structure associated with homo-oligomerization. The largest cluster involved intermolecular coiled coils (**Fig. 5A**), motivating an analysis of their representation across proteomes. While coiled coil regions can be detected from sequence alone ^35^, such predictions show higher false positive rates and lower sensitivity than structure-based assignment methods ^36–38^. Moreover, such sequence-based approaches cannot distinguish inter and intramolecular coiled coils, so our data uniquely enable comparing both types. Therefore, we used the structure-based method, SOCKET ^37^, to identify coiled coils in our quaternary structure models. Our analysis revealed that intra-molecular coiled coils exist at comparable frequencies across the four proteomes: 10.5% of proteins in the human dataset contained >5% of intramolecular coiled coils, versus 11% in yeast, 8.6% in *E. coli* and 7.8% in *P. furiosus* (**Fig. 5B**). In contrast, intermolecular coiled coils showed a marked increase in the human proteome, with 20.7% of homo-oligomers containing >5% of intermolecular coiled coils, versus 11.1%, 7.8%, and 4% for yeast, *E. coli*, and *P. furiosus*, respectively (**Fig. 5B**). This finding implies that coiled coil regions have been major enablers of quaternary structure evolution in the human lineage. Given the ability to design coiled coil interfaces ^39^, this finding opens prospects for designing coiled coil peptides and proteins to probe and intervene in many cellular processes.

**Figure 5.**
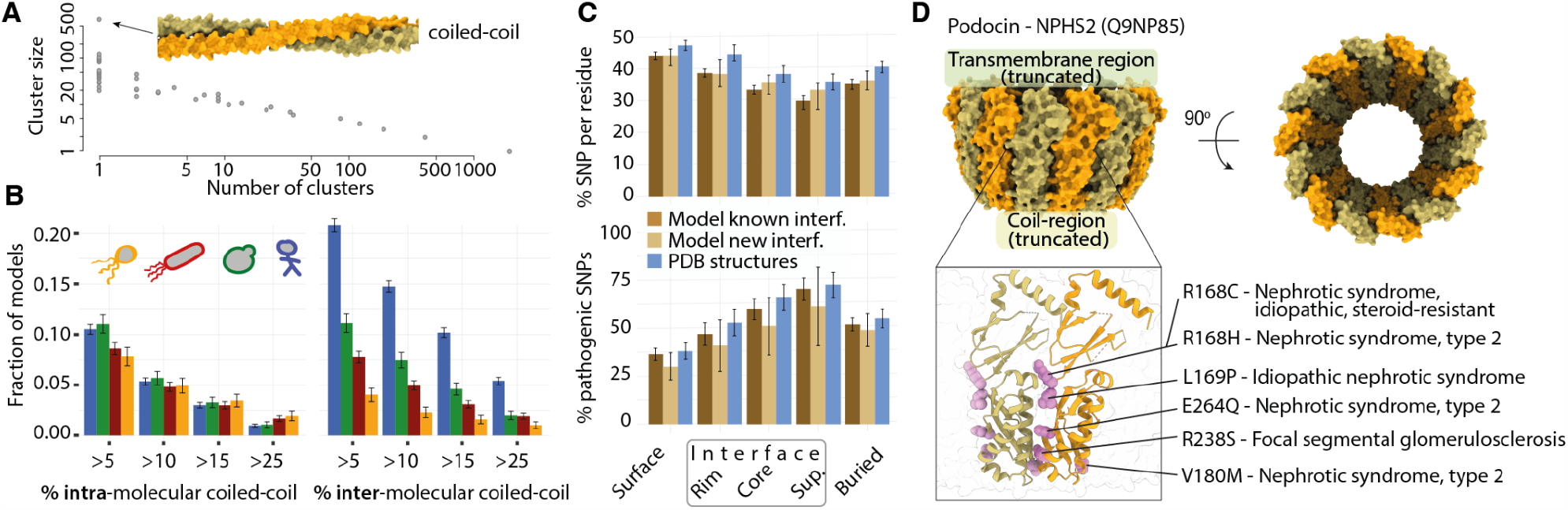
Analyzing the quaternary structure datasets provides insights into proteome evolution, disease mutations, and structure. **A**. Clustering all dimer models by structural similarity yields 2991 clusters. The scatter plot depicts the number of clusters having a particular size. At the extremes, there are 1957 singletons, and the largest cluster contains over 500 structures and corresponds to coiled coil-mediated dimerization. **B**. Barplot of the fraction of models containing intra (left) or intermolecular (right) coiled coils. Frequencies are comparable across kingdoms for intramolecular coiled coils, whereas intermolecular coiled coils are more prominent in Eukaryotes. **C**. Barplot depicting the percentage of SNPs normalized by residues (top) or pathogenic SNPs normalized by benign and pathogenic (bottom) across protein regions as defined in Ref. ^40^. Three types of structures are compared: human structures from the PDB (blue), models with quaternary structure type previously observed (dark brown) or that are novel (light brown). Error bars show the 95% confidence interval on the median (top) or mean (bottom). **D**. Predicted ring structure of podocin, which contains fourteen subunits and is bound to the plasma membrane through an alpha helix absent from the model. Two Podocin chains are highlighted, and interface residues with known pathogenic mutations from ClinVar ^41^ are shown (purple) along with the name of the associated condition.

Second, we projected 662,413 non-synonymous SNPs (single nucleotide polymorphisms) onto quaternary structures of the human dataset (Methods). We considered separately known structures and models, and further separated the models into those involving a known interface type or a novel one. We found that interfaces contained lower SNP frequencies than non-interface regions with equivalent solvent exposure (**Fig. 5C, Fig. S8**), consistent with the former being under stronger purifying selection than the latter (interface-core vs surface: *p-value*<0.0001; other comparisons and effect sizes are detailed in **Table S1**). Furthermore, interfaces showed significant enrichment in disease-associated SNPs when compared to non-interface regions (interface-core vs surface: +71.6%, p-value=0.0027; Table S1). The enrichment of disease-associated SNPs at interaction interfaces generalizes previous observations based on existing structures ^42^. Importantly, the enrichment is of a similar magnitude among experimentally characterized structures (+74.4%) supporting that the newly predicted quaternary structure types are as likely to be involved in diseases via interface mutations. A notable example is Podocin, a protein expressed in podocyte cells, which act as filters in the blood-urine barrier. Several mutations associated with nephrotic syndromes and renal failure appear at the interface of Podocin (Figure 5D), suggesting they impair its assembly into rings.

Third, we estimated the prevalence of symmetry among all protein complexes characterized to date by proteomics experiments. We gathered information on protein complex composition from multiple sources (Methods) and assigned each complex to one of four categories: symmetric homo-oligomer, symmetric heteromer, pseudo-symmetric heteromer, or asymmetric heteromer. Each assignment was made depending on the complex composition in homo-oligomer-forming proteins and paralogous sequences (Methods). We found that a majority of protein complexes form symmetrical assemblies (**Fig. 6, Data S4**). This is especially striking in *E. coli*, where more than 90% of the complexes form symmetrical homo or hetero-oligomers. In eukaryotes, we found that 60-65% of complexes are symmetric and these numbers increase to 65-70% when including pseudo-symmetries. These numbers highlight the ubiquity of symmetry in proteomes, which is key to consider when analyzing protein complex evolution ^43^ and assembly ^44,45^. Equally important, symmetry introduces multivalence, a fundamental property mediating assembly at the mesoscale by phase separation ^4^.

**Figure 6.**
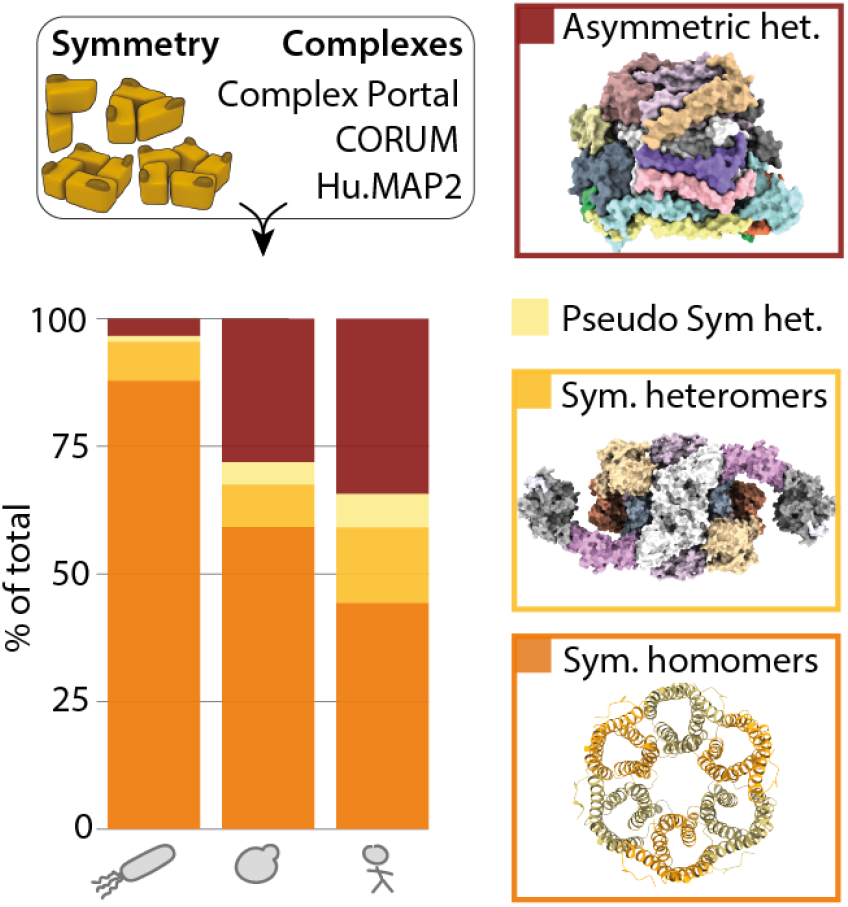
Integrating our datasets with omics data reveals a ubiquity of symmetry in cellular protein complexes. The proteome-wide quaternary structure information was integrated with omics data on protein complexes. Each complex (left) was assigned one of four types based on its protein composition: homo-oligomeric, symmetric heteromer, pseudo-symmetric heteromer, or asymmetric (Methods). This analysis reveals a striking ubiquity of symmetry among protein complexes.

## Conclusions

We have characterized protein quaternary structures across four proteomes with very high accuracy. Importantly, inferring quaternary structure information is challenging even when an experimental structure is characterized by X-ray crystallography. This is due to the difficulty in distinguishing fortuitous crystal contacts from physiological ones. These difficulties mean that upwards of 10% of biological assemblies available in the Protein Data Bank are estimated to be non-physiological ^46^. Consequently, the accuracy of our predictions can be compared to quaternary structure information originating from X-ray crystallography. Additionally, a significant advantage of our approach is the inclusion of full-length proteins, which enables capturing homo-oligomerization modes absent or altered in truncated proteins. The strategy devised in this work can be readily scaled up to cover a larger number of organisms relevant to fundamental and applied biology.

Here, we have focused on four proteomes - one archaeon, one bacteria, and two eukaryotes - and increased the quaternary structure coverage of these organisms by 50-100%. Notably, only a handful of new folds have been discovered among ∼600,000 monomeric structures predicted by AlphaFold2 ^47^. By contrast, we identify hundreds of novel quaternary structure types, implying that this space is much larger than that of tertiary structures. Remarkably, some of these novel quaternary structure types are conserved across the tree of life, provocatively highlighting their functional importance.

These proteome-wide datasets will aid biologists in gaining insights into specific proteins, and excitingly, it will provide a basis for a structure-guided understanding of proteome assembly. Here, we use these data to conduct three general analyses of proteomes’ quaternary structures. We observed that coiled coils are important mediators of quaternary structure evolution in eukaryotes. We found that interaction interfaces in homo-oligomers are hot-spots of disease-associated polymorphisms. Finally, we revealed that 60% of known protein complexes (both homo- and hetero-oligomeric) are symmetric in yeast and human, and this number increased to 90% in E. coli.

By expanding our knowledge of the protein quaternary structure space, this research opens up exciting new possibilities for interpreting network, omics, disease, and evolutionary data through a structural lens, ultimately aiding our understanding of the fundamental principles of proteome assembly and evolution.

## Methods

### Generation of homodimer models and monomers for human

The proteome sequences were downloaded from Uniprot for *P. furiosus* (proteome id: UP000001013), *E. coli* strain O157:H7 (proteome id UP000000558), *S. cerevisiae* (proteome id: UP000002311, reviewed entries) and *H. sapiens* (proteome id UP000005640, reviewed entries). The sequences of the proteomes were submitted to AlphaFold2 using a local implementation of ColabFold ^48^. MSAs were generated with the MMseqs2 online service ^49^ and saved locally for later reuse. Sequences longer than 1200 amino acids were not processed as the effective length of dimers for such proteins exceeds 2400 amino acids, for which predictions became too GPU and memory expensive. Five models were generated for each protein, and information relating to these structures was stored in a MySQL relational database. The same protocol and saved MSAs were used to generate monomeric structural models of the *H. sapiens* proteome.

### Structure processing: contacts, clashes, and core structure definition

Intra- and intermolecular residue contacts were identified using the same definitions as in 3DComplex ^50^. We quantified clashes in two ways: the first was atom pairs closer than 2 Å. In addition, we calculated a clashscore using the software Phenix ^51^. Solvent accessibility and interface sizes were determined using FreeSASA ^52^. The residues stabilized in the dimer structure (Figure 2E) are defined as those exhibiting at least two inter-chain contacts and at most one intra-chain contact. All residues were considered for inter-chain contacts, and residues from *i*-4 to *i*+4 were excluded in calculating intra-chain contacts of residue *i*.

Structural models are computed on full-length protein sequences. As a result, they often contain flexible, disordered regions. Several analyses require comparing structures, and the presence of such flexible regions or domains can mask an otherwise conserved core. We used a three steps procedure (**Fig. S1A**) to trim these regions and define the “core structure”: (i) we discarded residues with a pLDDT score below 40; (ii) of the remaining residues, a median pLDDT score was computed, and residues with a pLDDT score below 75 and below the median value are discarded; (iii) we applied single linkage clustering on the contact matrix of the remaining residues and retained the largest cluster, thus eliminating disconnected structural parts.

### Selecting a representative model per protein

Out of the five models generated for each protein sequence, we selected one representative per protein. We performed a pairwise structural superposition of the models’ structure using Kpax ^53^. Structures were matched when they showed a TM-score above 0.75. We note that these matches are not necessarily reciprocal because structures can have different lengths after trimming flexible regions. Within a group, the structure with the highest dimer probability (pae4-con3 metric, defined later) was kept as a reference. Structures showing more than 10% of atomic clashes at the interface, or more than 10% of residues with clashes were discarded. If all models in a group exceeded these clash thresholds, the selected representative structure was the one with the fewest clashes at the interface.

### Evaluating the confidence of homodimer models

First, we assembled a dataset to assess the power of specific metrics returned by AlphaFold2 (e.g., inter-chain PAE) or computed on the models (e.g., size of the interface) to discriminate between monomers and dimers. The dataset was derived from the PDB ^17^ and consisted of 349 monomers and 77 homodimers non-redundant at a level of 30% sequence identity and elucidated by X-ray crystallography. The structures in the dataset were deposited after May 2018, were not part of the AlphaFold2 training set, and no structure prior to that date showed >35 % identity and >50% overlap. As crystallographic structures can contain fortuitous interactions mediating the crystal assembly, we excluded low-confidence monomers and homodimers (error probability >25%), as evaluated by QSBIO ^54^. We computed AlphaFold2 predictions for these structures, using the corresponding Uniprot ID and sequence. Since our proteome predictions are performed on full-length proteins, we also employed the full-length proteins in this benchmark. We derived several metrics on the predicted model structures, and all were highly informative for discriminating between correct homodimer models (TM-score > 0.8 with experimental structure) and monomers in the dataset (**Fig. 1A, Fig. S1B**).

We used the following metrics individually:

1. The number of residue-residue contacts in the core structures (metric identified as “*con3”*);
2. The mean inter-chain PAE values of interface residues in the core structure (metric identified as “*pae3”*);
3. The mean PAE values of contact (i.e., pixels) in the core structures (metric identified as “*pae4”*);
4. The consistency of the five models as determined from the structural superposition of their core structure before the single linkage clustering was applied. Each model was assigned a score ranging from 0 (where no other model was structurally similar) to 8 when a model reciprocally matched all the other four models (metric identified as “*repre2”*). The name in parentheses is used to refer to each metric later in the text.

We converted these four metrics into probabilities by fitting a logistic regression model based on the benchmark data, and also added the combination of the *pae4* and *con3* metrics, henceforth *pae4con3*. The logistic regression models subsequently enabled scoring any dimer model to yield a probability ranging between 0 (likely monomer) and 1 (likely dimer).

Five probabilities were calculated for the 156,065 models. As each metric showed a strong discriminatory power, we were inclusive and initially retained all models with any of the five probabilities above 0.5. We did not consider structures where the number of interface clashes represented more than 10% of the contacts, where the number of residues involving clashes represented more than 5% of the protein length, or where the number of contacts in the core structure or full-length protein was less than 10 or 30 respectively.

### Identifying structures related to the four proteomes in the PDB

The sequences of the four reference proteomes were searched against known structures released up until March 1st 2022. We used FASTA ^55^ as well as HHblits ^56^ to perform pairwise alignments and stored information about the hits, their corresponding sequence identity, and sequence coverage in two separate MySQL tables. Each model was assigned to one of three categories of sequence identity: 0-30%, 30-70%, and 70-100%. We imposed an overlap of at least 50% of the core structure to accept a match. We estimate the increase in structural coverage of a proteome as the number of proteins in the first category divided by the number in the third.

### Clustering quaternary structures from the reference set

The core structure of each dimer model from the datasets was structurally supposed with KPAX ^53^ to all other models sharing one or more PFAM domains ^29^. A binary matrix was derived where all structure pairs showing a TM-score above 0.7 were set to 1, and the rest to 0. Single-linkage clustering was applied to this matrix, yielding 2991 quaternary structure distinct clusters or “types”.

### Defining novel quaternary structure types

Dimer models were superposed to all putative homologs identified with FASTA and HHBLITS. We employed two superposition methods: KPAX ^53^ and MMalign ^57^ and kept the highest TM-score. We initially excluded as novel any dimer model for which a TM-score higher than 0.65 was identified. We note that this cut-off has to be different from the threshold classically used for monomers (0.5) because the same protein forming two distinct dimer structures will show a minimum TM-score of 0.5. We previously found the value of 0.65 to be optimal ^54^. Additionally, we imposed that at least 50% of residues in each chain of the dimer core structure should be matched. Finally, we only considered a quaternary structure type as novel when all those in the same cluster were novel themselves.

### Symmetry detection, assignment, and reconstruction

We used AnAnaS ^25^ to identify symmetries in the homodimer models. We first assigned C2 symmetry to a model when the RMSD calculated by AnAnaS was below 4Å and the clash score below 200. When a C2 symmetry was not detected, we searched for higher-order symmetries, from C3 to C12. The symmetry with the lowest RMSD was then retained, provided it showed a value below 4 Å for C3, 3.5 Å for C4, 3 Å for C5, and 2.5 Å for C6-C12, all with a clash score below 200. If the best symmetry detected was C12, higher symmetries were searched, from C13 to C24. The symmetry with the best RMSD was retained, providing it had a better RMSD than the C12 symmetry, and an RMSD below 2 Å for C13-14, and 1.5 Å for C15-23 with a clash score below 200.

At this stage, structures with no detected symmetry encompass monomers, dimers where both subunits are flexible and where a symmetry axis cannot be reliably defined, and proteins that form infinite assemblies, such as actin filaments.

To distinguish between those three types, we relied on the symmetry of the contact matrix that enables distinguishing homotypic from heterotypic interactions. We listed all residue pairs exhibiting more than one atom in contact (e.g., number 10 in chain A with number 40 in chain B), and recorded the fraction of those showing a reciprocal contact (i.e., number 40 in chain A with number 10 in chain B), including reciprocal pairs with a single atomic contact. If all residues in contact are reciprocal, the interface is necessarily homotypic. However, C2 symmetry may not be detected due to structural flexibility. Conversely, a homotypic score of 0 means the dimer is compatible with helical or cyclic symmetry. Thus, we combined this residue-based symmetry information with global symmetry information and defined three additional categories:

- *Trans*: dimers that show no global symmetry (identified with AnAnaS) and a homotypic score of 0, which implies the formation of filaments with translational or helical symmetry.
- *C2-flex:* dimers with no global symmetry but pronounced local symmetry (homotypic score >= 0.4). These typically result from flexible structures with local C2 symmetry axes that are not aligned globally across the protein.
- *Ambiguous:* dimers with no global symmetry and limited local symmetry (homotypic score >0 and <0.4). In these structures, different regions can display incompatible symmetries, e.g., one domain exhibiting C2 symmetry and another adopting a translational symmetry.

### Comparing monomer and homodimer human models

To assess the structural similarity between the monomers and dimers predicted by AlphaFold2, the core structure of the 5 monomeric models was superposed onto both subunits of the corresponding dimer core structure from the reference set using TMalign ^58^. The core structure of the dimer was trimmed to remove residues not present in the core structure of the monomer (in order to not artificially decrease the TM score). The TM-scores were normalized by the length of the chain of the dimer and the final TM-score was the highest of the 10 scores.

### Comparing the models against recently solved cryo-EM structures

We selected 23 structures of human proteins solved by electron microscopy, released after May 2018, and for which no structure released prior to that date shared >35% sequence identity with a sequence coverage of 50% or more. After manual curation, three structures were discarded from the set because the quaternary structure state was ambiguous (detailed in **Fig. S4** and **Data S3**). The core models were further trimmed to remove residues not present in the EM structure and were superposed using MMalign ^57^ or TMalign ^58^ to superpose monomers.

### Detection of coiled coil domains

Coiled coils were detected structurally using the software SOCKET2 ^37^, with a packing cutoff of 7 Å. This identifies the knobs-into-holes packing signature of coiled coil structures rather than sequence-based signatures such as heptad repeats ^36^. Coiled coils were classified into two categories: intra and intermolecular coiled coils, which are defined as those involving residues belonging to the same or different chains, respectively.

### Generation of final models

All dimer models were relaxed using openMM v.7.3.1 ^59^ in a constrained AMBER forcefield ^59^, identical to the protocol previously published with AlphaFold2 ^8^. We provide the dimer structures of all models, with both original and relaxed coordinates. Ring complexes were generated with AnAnaS ^25^ based on the dimer core-structure. Importantly, flexible segments absent from the core-structure could occupy the space of a ring subunit adjacent to the dimer, leading to extensive steric clashes in the full-length symmetry-based reconstruction. Thus, to include the flexible segments while eliminating steric clashes, the assembly had to be repredicted in the context of the complete ring. However, such predictions are very demanding both GPU and memory-wise, and we also noticed that AlphaFold2 converged less efficiently for large complexes. For these reasons, we developed a protocol tailored to reconstruct large assemblies given a known ring template with missing segments. First, the AnAnaS ^25^ reconstructed model containing only amino-acids defined in the core-structure was used as a template input to AlphaFold2 to facilitate its convergence while avoiding the need for memory heavy multiple sequence alignments. To give the AlphaFold2 predictions more flexibility, we masked all sequence information in the template by changing the sequence to all gap-tokens and masking out all sidechain atoms except for C-beta, given that the C-beta distance matrix is one of the inputs. In the case of glycine, a virtual C-beta atom was added to the template. However, the sole use of a template was often not sufficient to enable AlphaFold2 to predict the correct structure, and we overcame this issue by initializing the coordinates using the template. We call this protocol “big bang” initialization, which contrasts with the default where coordinates are initialized at zero. We therefore used the template to initialize the backbone coordinates for the first iteration in the structure module. In the case of discontinuous segments in the template, we initialized the coordinates as an interpolation between the start and end residue of the missing segment. After this process, the final models were aligned with the initial ring (predicted based on core-dimer symmetry) using MMalign ^57^ to ensure they were similar. Only four structures out of 955 displayed a TM-score < 0.5 and these were discarded from the final set.

### Analysis of SNPs

Protein sequences were aligned to genomic locations using Ensembl Variant Predictor (VEP) ^60^. Human SNP data were obtained from gnomeAD v2.1.1 mapped to GRCh38 and matched to the protein sequence ^61^. Information on disease-associated SNPs was extracted from ClinVar (version 20230115) ^62^. We counted as disease SNPs those that contained the strings ‘pathogen’ or ‘risk’ in their ClinVar description. Benign SNPs were those containing the string ‘benign’. Other descriptions were discarded.

We analyzed the frequency of SNPs and pathogenic SNPs in different protein structural regions defined in ^40^. The SNP frequency in a region was calculated per protein as the number of SNPs in the region divided by the number of residues. The distributions of these frequencies per region are described in Figure S5 and Figure 4B summarizes the median of each distribution along with a 95% confidence interval estimated from 10,000 bootstraps. When comparing regions we computed p-values under a null hypothesis where medians were equal. The p-values were inferred from the bootstrap data, as the fraction of iterations where the median value for the interface rim|core|support region was greater than the median for the surface|surface|interior respectively (Table S1). For disease-associated SNPs we followed the same process, except that we focused on the mean of the distributions because they included mostly discrete values (0/1) due to the small numbers of pathogenic and benign SNPs. The distribution of pathogenic SNP frequencies is shown in Figure S5B, and Figure 4B summarizes the mean of the distributions along with a 95% confidence interval.

### Prevalence of symmetry in protein complexes including heteromers

Analyzing the prevalence of symmetry at the complexome level required comprehensive information on protein complexes. We therefore identified protein complexes using several databases:

- *H. sapiens* complexes were collected from Humap2 ^63^ using high-confidence entries (confidence score <= 3), CORUM ^64^ and Complex Portal ^65^.
- *S. cerevisiae* complexes were collected from YeastCyc ^66^, CYC2008 ^67^, YHTP2008 ^67^ and Complex Portal ^65^.
- *E. coli* complexes were collected from EcoCyc ^68,69^ and Complex Portal ^65^.
- *P. furiosus* was not analyzed due to the lack of information on complexes.

For each organism, we concatenated information from the databases and then removed redundancies. two complexes sharing at least 80% of subunits (identified through UNIPROT identifiers) were considered redundant, and only the largest was kept. These filters resulted in 3489, 590 and 265 non-redundant complexes for *H. sapiens, S. cerevisiae* and *E. coli* involving respectively 6217, 2149, and 759 unique proteins. Combining these data with our datasets gave 6234, 1445, and 2167 homo or hetero complexes corresponding to 8511, 2957 and 2665 proteins, respectively.

Next, we categorized each complex according to its composition in homo-oligomer-forming proteins, and based on the presence of paralogous chains. Two proteins were considered paralogs if they showed any degree of sequence similarity and an alignment overlap of at least 60%, or shared the same PFAM domain architecture. Based on this information four categories were defined:

- Symmetric homomers contain a single subunit.
- Symmetric heterocomplexes contain over 30% of homo-oligomerizing subunits and none of them has a paralogous chain in the complex.
- Pseudo-symmetric heterocomplexes contain over 30% of homo-oligomerizing subunits and at least one of them has a paralog in the complex.
- Asymmetrical heterocomplexes contain less than 30% of homo-oligomerizing subunits.

### Q8U0N8 protein purification

The Q8U0N8 gene from *Pyrococcus furiosus* has been ordered from Twist Biosciences as a synthetic gene cloned into the pET21 vector with C-terminal His_6_-tag. The protein was expressed in *E. coli* BL21 DE3 cells and expressed overnight with 0.5 mM IPTG at 18 °C in LB medium supplemented with 50 μg/ml ampicillin. The pellet was resuspended and sonicated in 20 mM HEPES pH 7.5, 500 mM NaCl, 50 mM L-arginine, 10% glycerol buffer supplemented with 1 mM PMSF, and 125 μg/ml DNase. Cell lysates were clarified using ultracentrifugation and loaded on a 5 ml Ni-NTA Superflow column (QIAGEN) and washed with 20 mM HEPES pH 7.5, 500 mM NaCl, 50 mM L-arginine buffer with imidazole ranging from 10-30 mM, and subsequently eluted with 300 mM imidazole. Main protein fractions were concentrated and injected onto a Superose S6 10/300 gel filtration column (GE Healthcare) in 20 mM HEPES pH 7.5, 300 mM NaCl. Protein fractions were concentrated, flash-frozen in liquid nitrogen, and stored at -80 °C.

### Molecular weight determination by mass photometry

Mass photometry experiments were conducted using a Refeyn TwoMP system (Refeyn Ltd., Oxford, UK) equipped with the AcquireMP and DiscoverMP software packages for data acquisition and analysis, respectively, utilizing standard settings. For the experiments, microscope coverslips of high precision, sourced from Refeyn, were utilized on a one-time basis. To maintain the droplet shape of the sample, self-adhesive silicone culture wells (Grace Bio-Labs reusable CultureWell™ gaskets) were employed. For contrast-to-mass calibration, Bovine Serum Albumin Fraction V low Heavy Metals (Millipore) oligomers with molecular weights of 66, 132, 198, and 264 kDa were employed. Prior to the measurements, protein stocks were diluted in stock buffers containing 20 mM HEPES pH 7.5 and 300 mM NaCl. Specifically, 2 µL of the protein solution was combined with 18 µL of analysis buffer, resulting in a final drop volume of 20 µL with a concentration of approximately 1 µg/mL.

### cryoEM sample preparation, data collection, processing, and structure building

The Q8U0N8 hexameric complex was diluted to a concentration of 4 mg/ml. 4 µl were applied to a glow discharged 300-mesh holey carbon grid (Au 1.2/1.3 Quantifoil Micro Tools), blotted for 1.5–2.5 s at 95% humidity, 10 °C, plunge frozen in liquid ethane (Vitrobot, FEI) and stored in liquid nitrogen. Data collection was performed on a 300 kV FEI Titan Krios G4 microscope equipped with a FEI Falcon IV detector. Micrographs were recorded at a calibrated magnification of 168,674× with a pixel size of 0.83 Å and a nominal defocus ranging from −0.8 μm to −2 μm.

Acquired cryo-EM data was processed using cryoSPARC ^70^. Gain-corrected micrographs were imported, and micrographs with a resolution estimation worse than 5 Å were discarded after patch CTF estimation.

Initial particles were picked using blob picker with 100–140 Å particle size. Particles were extracted with a box size of 360 × 360 pixels, down-sampled to 120 × 120 pixels. After 2D classification, clean particles were used for *ab initio* 3D reconstruction. After several rounds of 3D classification, the class with most detailed features was re-extracted using full 360 × 360 pixel box size and subjected to non-uniform and local refinement to generate high-resolution reconstructions ^71^. The local resolution was calculated and visualized using ChimeraX ^72^.

For structure building, the *in silico* model generated in our study was split into segments and docked into density using ChimeraX. Subsequent manual model adjustment and refinement was completed using COOT ^73^. Atomic model refinement was performed using Phenix.real_space_refine ^74^. The quality of the refined model was assessed using MolProbity ^51^. Structural figures were generated using ChimeraX. The refined atomic model and corresponding cryoEM map was deposited under PDB: 8P49 and EMDB: 17402, respectively.

## Acknowledgments

We thank Sergei Grudinin for help with AnAnaS, Harry Greenblatt, and the HPC Weizmann team for help with the computer infrastructure. We thank the Protein Production and Structure Characterization Core Facility, EPFL, Switzerland for help with protein purification. We thank the Dubochet Center for Imaging (DCI), EPFL-UNIL-UNIGE, Switzerland for cryoEM data collection. We thank Moran Shalev Benami, Patrick Barth, Anil Pasupulati, Sven Dahms, Robert Jefferson, and the members of EDL and BEC laboratories for helpful discussions.

## Authors contributions

HS, SD, EDL carried out structure processing and analyses; TL carried out the SNP analysis with help from BD; PK and HS did the coiled-coil analysis, CAG refined the structures and generated the final models; MP, YD, LJD, SG carried out protein expression assays; MP processed cryoEM data and structures; DW, BEC, SD, EDL oversaw and funded the research. SO provided computational protocols for structure refinement. HS and EDL wrote the manuscript with input from all authors.

## Data availability

We provide the data as Data Tables S1-3 as well as all structures generated, which are available on FigShare (https://figshare.com/s/af3c1d5969f7468f2caa). The protocol for AlphaFold2 “big bang” is accessible at https://github.com/sokrypton/ColabDesign.

