## Supplementary Material for "An atlas of protein homo-oligomerization across domains of life"

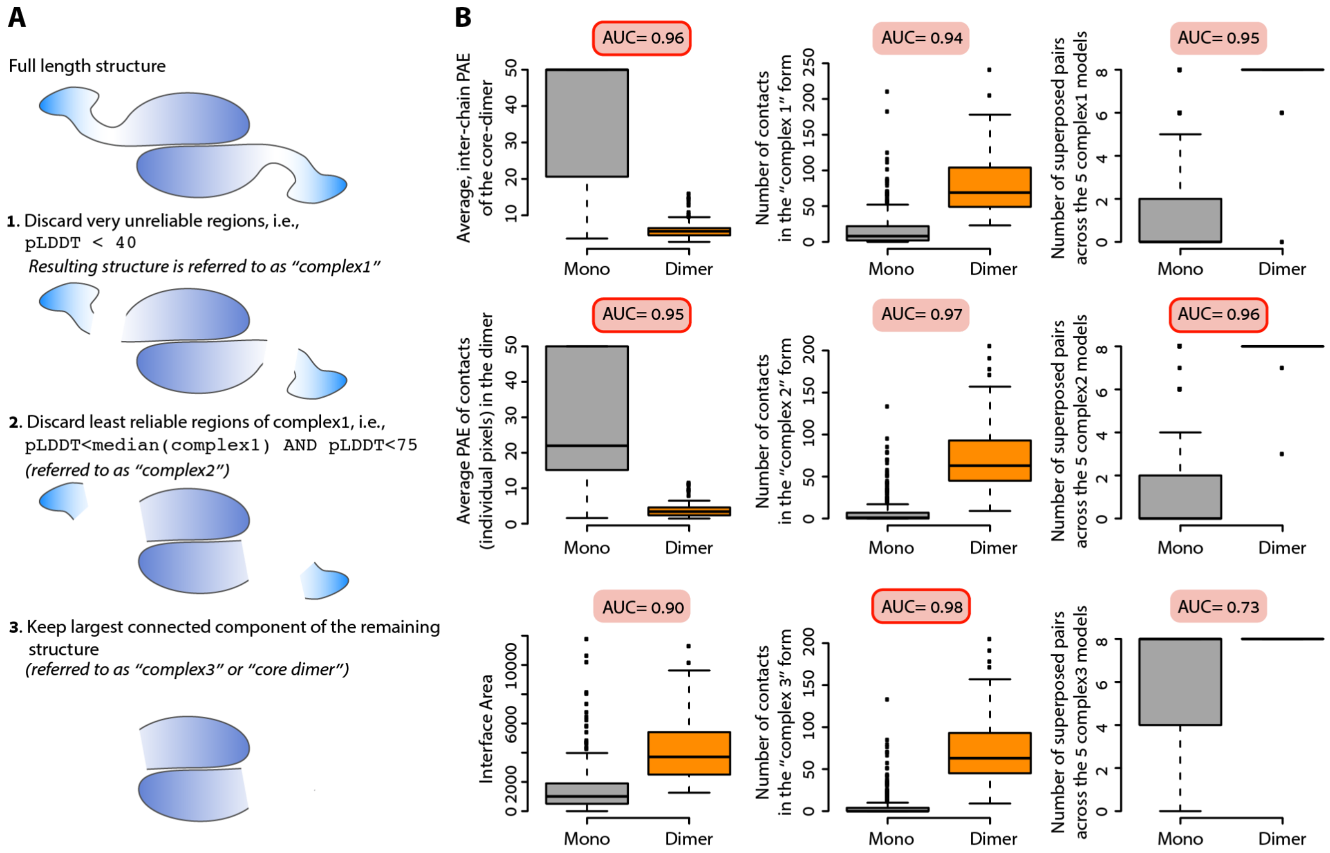


**Supplementary Figure 1. Structure processing to remove low confidence regions and accuracy of several metrics to discriminate monomer and dimer states from the predicted structure models.**
**A**. Structure models predicted by AlphaFold2 frequently contain flexible regions. When comparing structures, these regions artificially lower the structural similarity due to their conformational flexibility. Therefore, several of the analyses in this work focused on structures where these regions were truncated. The procedure followed to truncate these regions is highlighted. First, all residues with a pLDDT value below 40 were discarded. Second, we calculate the median pLDDT score of the remaining residues and discard those with a pLDDT below this median score provided their value is not above 75. Third, we discarded residues and segments disconnected from the core structure, with the latter being defined as the largest connected component in the residue contact matrix.
**B**. We analyzed several features for their ability to discriminate homodimers (orange) from monomers (grey) in our benchmark dataset. Several of the metrics that we tested proved reliable. Four metrics were eventually used to derive the reference sets of quaternary structures: the average inter-chain position alignment error (PAE) of the residues in contact (referred to as *pae3*), the average interchain PAE of the individual contacts (referred to as *pae4*), the number of contacts at the interface of the core structure (referred to as *con3*), and the number of superposed pairs across the five models generated by AlphaFold2, considering the truncated "complex2" structures (panel A).

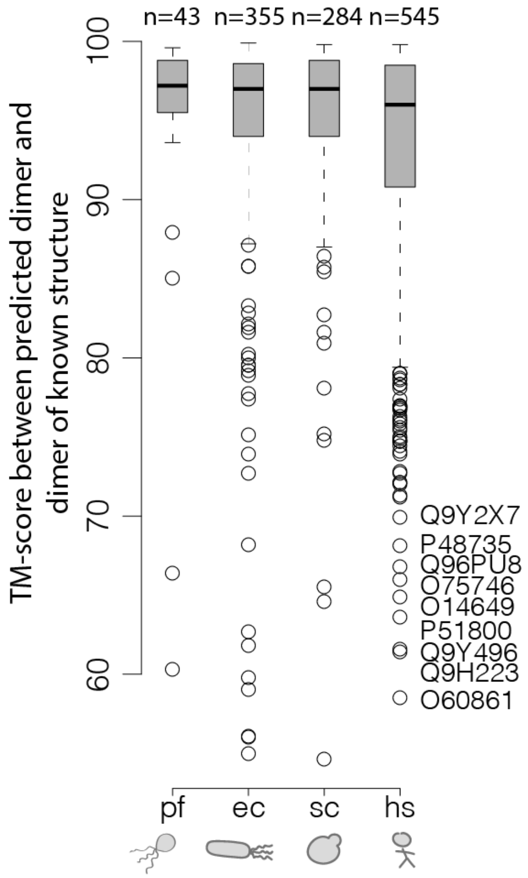


**Supplementary Figure 2. Dimers models show close agreement with their matching experimental structure.** The dimer structures from our dataset were compared to those of closely related structures in the PDB (sequence identity > 90%) for which the QSBIO error probability was below 25% [1]. The distribution of TM-scores of dimer structure pairs is shown for each species. In the human dataset, only seven structures exhibit a TM-score below 0.7, and we provide the matching UNIPROT identifiers. Overall, these discrepancies are not caused by incorrect predictions. Instead, they originate in conformational flexibility of the monomers (7 cases) or in experimental artifacts (2 cases), as detailed below:

1. 4dnn and Q96PU8. The low TM-score is due to 4dnn containing a large number of selenomethionine residues, which were not counted in the structural superposition. Inspection of the two structures reveals an excellent agreement between the two interface geometries.
2. 5mtv and Q9H223. The low TM-score is due to the conformational flexibility of each monomer, but the dimeric interface is highly similar between the PDB structure and the dimer model from our reference set.
3. 4p5x and O75746. The low TM-score is due to conformational flexibility of the monomers, while the interaction geometry is similar.
4. 2w6a and Q9Y2X7. The low TM-score is due to the core-structure and the X-ray structure not showing a large enough overlap. The core-structure of the model is shorter, but the part present in both the PDB and the core-structure interacts in the same manner.
5. 6iko and O60861. The low TM-score is due to conformational flexibility of each monomer, and the dimeric interface is highly similar between the PDB structure and the dimer model.
6. 2pfi and P51800. The low TM-score is due to a poor overlap between regions seen in the PDB structure and regions present in the core-structure. However, the interface region is highly similar.
7. 5jx1 and Q9Y496. The low structural similarity is due to 5jx1 being a chimeric protein with only a small region matching Q9Y496.
8. 4ja8 and P48735. The low TM-score is due to conformational flexibility of each monomer, and the dimeric interface is highly similar between the PDB structure and the dimer model.
9. 6rv2 and O14649. Both quaternary structures are very similar, but the TM-score is low due to a poor overlap with missing regions in the PDB structure and missing regions in the dimer core-model.

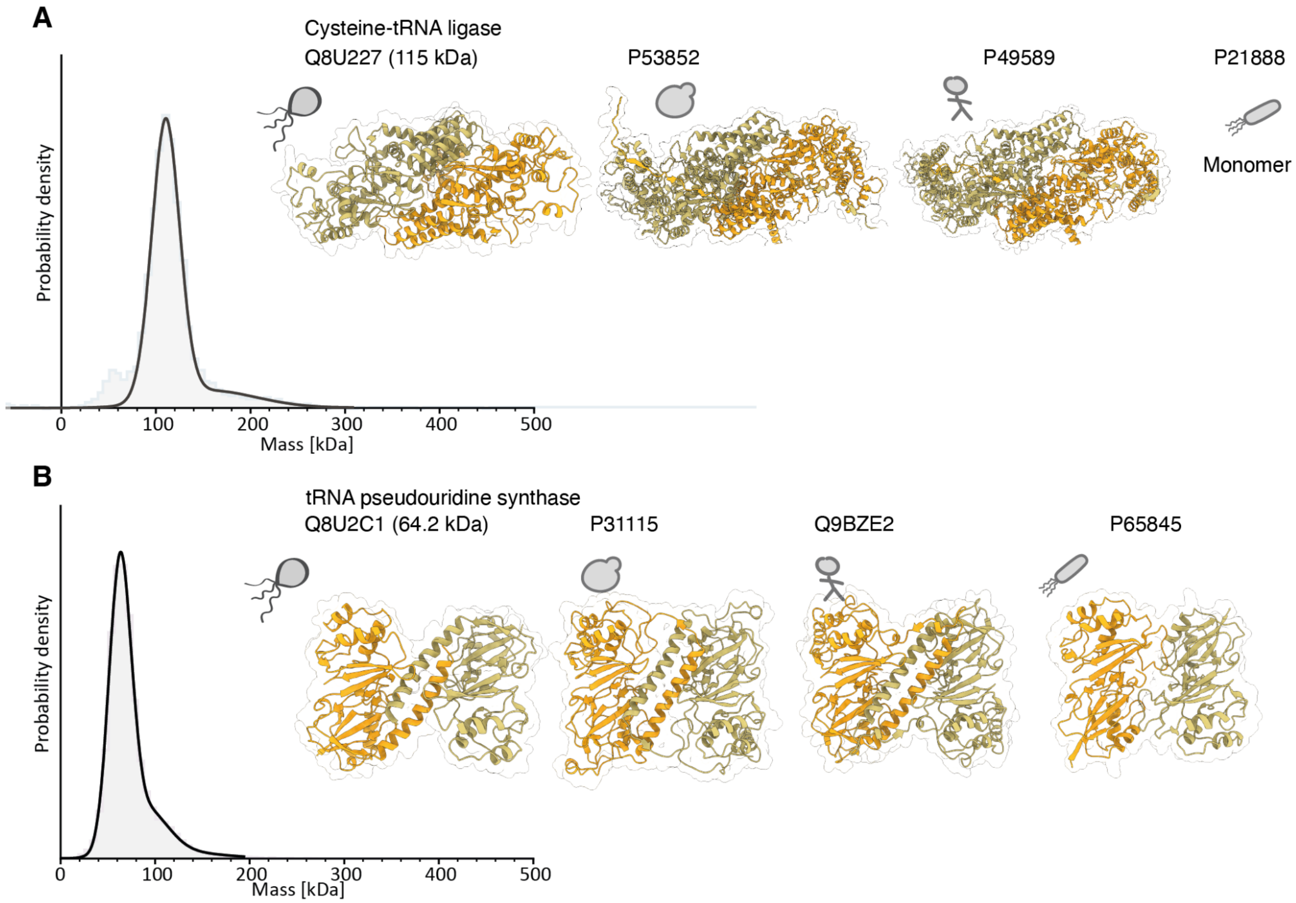


**Supplementary Figure 3. Experimental validation of the dimeric state corresponding to novel interface types.** **A**. Cysteine tRNA ligase from P. furiosus (Q8U227) was predicted as forming a homodimer. Similar dimers were also predicted in yeast (P53852) and human (P49589), and this dimer form was absent from PDB and represents a novel interface type. A crystal structure of a homolog from E. coli (P21888) is monomeric. The oligomeric state of the purified protein from P. furiosus appears dimeric, with a peak at 115 kDa. **B.** A tRNA pseudouridine synthase from P. furiosus (Q8U2C1) was predicted as forming a homodimer. Similar dimers were also predicted in yeast (P31115) and human (Q9BZE2). This dimer form was absent from PDB and represents a novel interface type. Interestingly, a crystal structure of a homolog from E. coli (P65845) also forms a dimer. Although the interaction geometry between the two chains is similar, the interface is entirely different. The oligomeric state of the purified protein from P. furiosus appears dimeric, with a peak at 69 kDa.

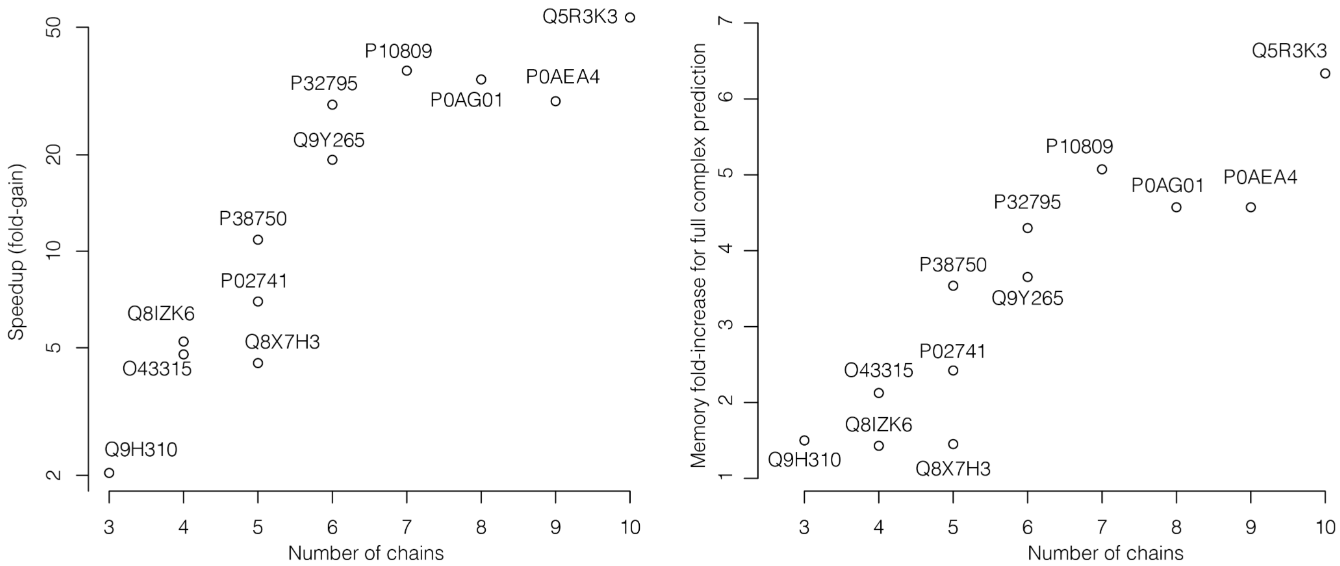


**Supplementary Figure 4. Time and memory requirements in predicting the structure of dimer models or the respective full-size complex.** Models and full complexes were predicted on the same machine and GPU (RTX6000 ADA, 48GB). The first three models were generated with three recycles each. The time and memory requirements were averaged for models 2 and 3, and the ratio of “complex / dimer” is displayed for each structure. We observe a 50-fold speedup for a complex with 10 subunits (left), and a >6-fold gain in memory requirements (right).

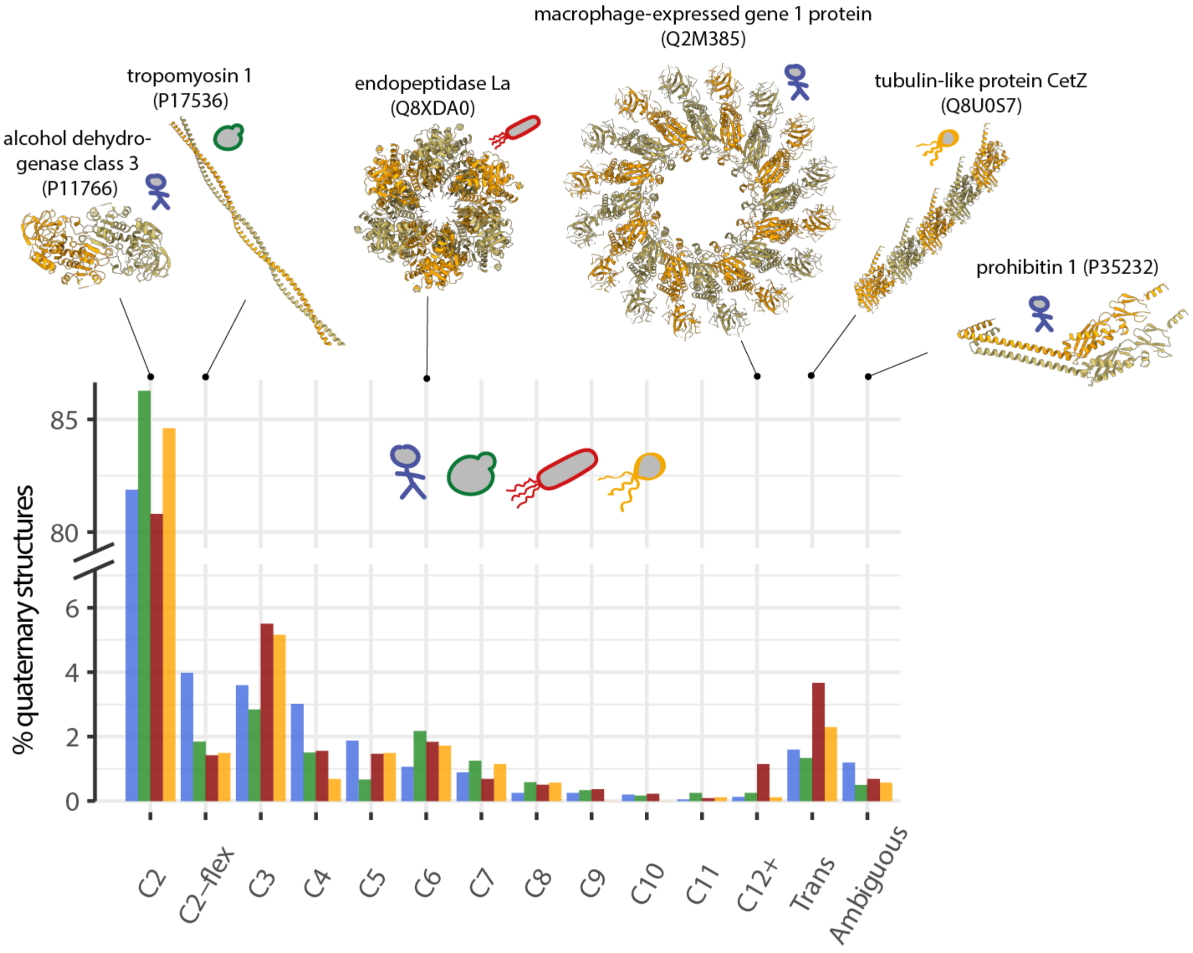


**Supplementary Figure 5. Distribution of homo-oligomer symmetry across organisms.** The percentage of each symmetry type is shown. Dimers represent the largest class (note the axis break). *Cn* represents point group cyclic symmetries and those involving 12 or more subunits were grouped in one class. The categories C2-flex, Trans, and Ambiguous are defined as described in the Methods section "Symmetry detection, assignment, and reconstruction". The alcohol dehydrogenase class 1 (UniProt id Q2M385) is an example of a C2 complex. The tropomyosin 1 (UniProt id P17536) is an example of a flexible C2 complex. The endopeptidase La (UniProt id Q8XDA0) is a C6 complex. The macrophage-expressed gene 1 protein (UniProt id Q2M385) is an example of a large cyclic complex comprising 16 subunits. The tubulin-like protein CetZ (UniProt id Q8U0S7) is an example of a complex showing a Trans symmetry susceptible to form infinite assemblies. The prohibitin 1 (UniProt id P35232) is an ambiguous case where no symmetry could be detected because of the flexible C-ter alpha helix forming a coiled-coil interface between the two chains of the model.

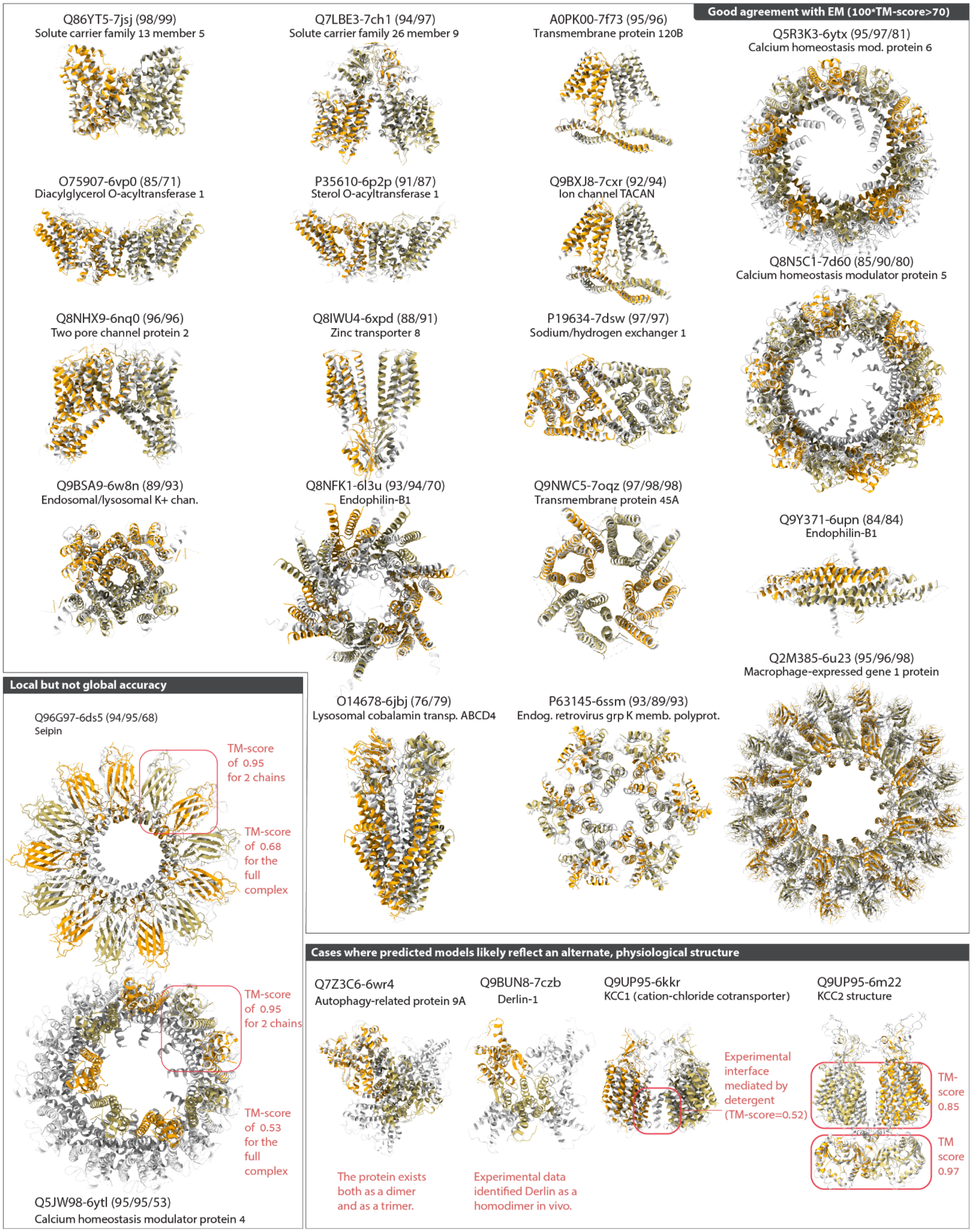


**Supplementary Figure 6. Quaternary structure models accurately recapitulate recent structures solved by electron microscopy.** We identified structures of human proteins solved by electron microscopy with no homolog in the Protein Data Bank prior to May 2018 (>35% sequence identity). We then superposed predicted models onto these structures. Each superposed pair is shown with the model in orange-green and the experimental structure in white. We provide the UniProt and PDB codes above each pair along with scores (TM-score x 100). The scores represent the structural similarity of the monomer, dimer, and full complex superposition. Most models agree with the experimental structure (score > 70). Two models show excellent chain-chain interaction geometry (dimer score = 95) but poor global structure similarity due to inconsistent numbers of chains. In three cases where the prediction and experimental structure differ significantly, we highlight the ambiguous nature of the experimental data. The structure identified as 6wr4 exists both as a dimer and trimer, as noted in the original publication [2]. Interestingly, the dimer structure that we model involves the same interfaces as those seen in the experimentally characterized trimer. Therefore, our model may capture the alternative dimer state observed experimentally. In a second example, the structure 7czb forms a tetramer while our model is dimeric. Derlin-1 has been observed to form homodimers in vivo [3] and was also observed to be part of the hetero-oligomeric Hrd1 ubiquitin ligase complex, within which it exists as a monomer [4]. These observations imply that it can adopt multiple oligomeric states, and our model may capture the experimentally observed homodimer state. In the last example, the structure 6kkr consists of the transmembrane domain of the cation-chloride co-transporter KCC1. The structure shows a dimeric assembly mediated by detergent molecules. Our model shows a different interaction mode where dimerization is mediated by the cytosolic domain (which is absent from the structure 6kkr). Interestingly, the assembly seen in our model is similar to that of a homolog characterized more recently (KCC3, PDB code 6m22). Such similarity supports the validity of our model and suggests that the detergent-mediated interface represents an alternative interaction mode or could result from using a truncated construct. Promiscuous or non-physiological protein-protein interfaces are highly evolvable [5] and are pervasive in X-ray crystallography experiments [6]. The lower protein concentrations required for cryo-electron microscopy experiments make such interactions unlikely, yet certain contexts (truncations, concentration in membranes) might nevertheless promote their formation.

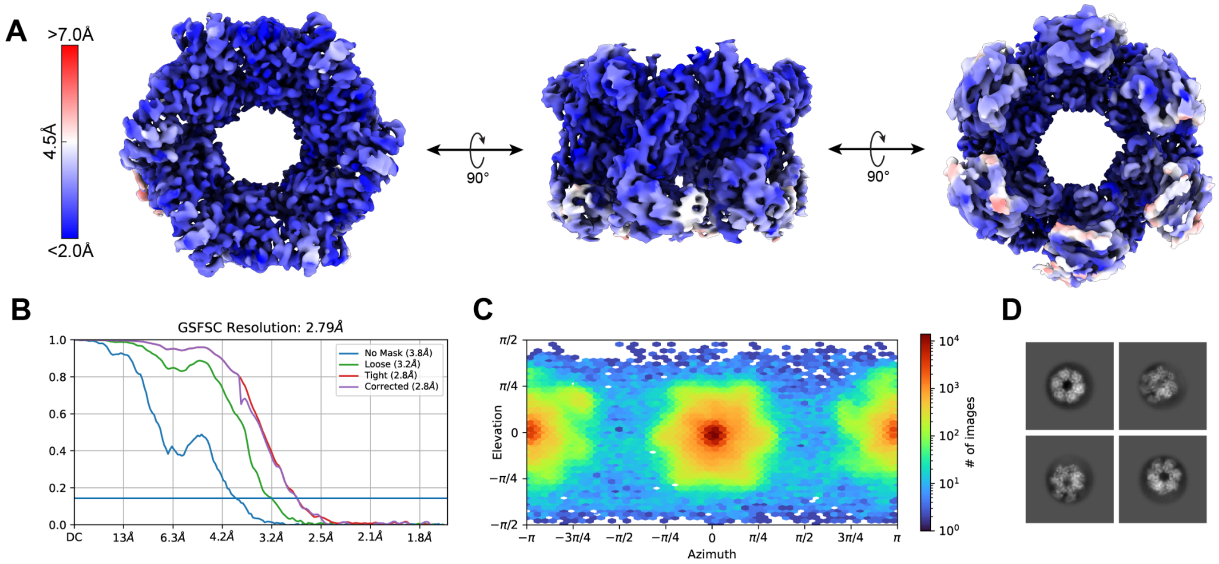


**Supplementary Figure 7. Cryo-EM processing and results for the Q8U0N8 hexamer. A.** Top (left), side (middle), and bottom (right) views of the unsharpened cryo-EM density map of the Q8U0N8 protein hexamer. The maps are colored by local resolution. **B.** Gold-standard FSC curve with resolution cut-off indicated at 0.143. **C.** Particle distribution heatmap of the final reconstruction. **D.** Example 2D class averages of different particle views.

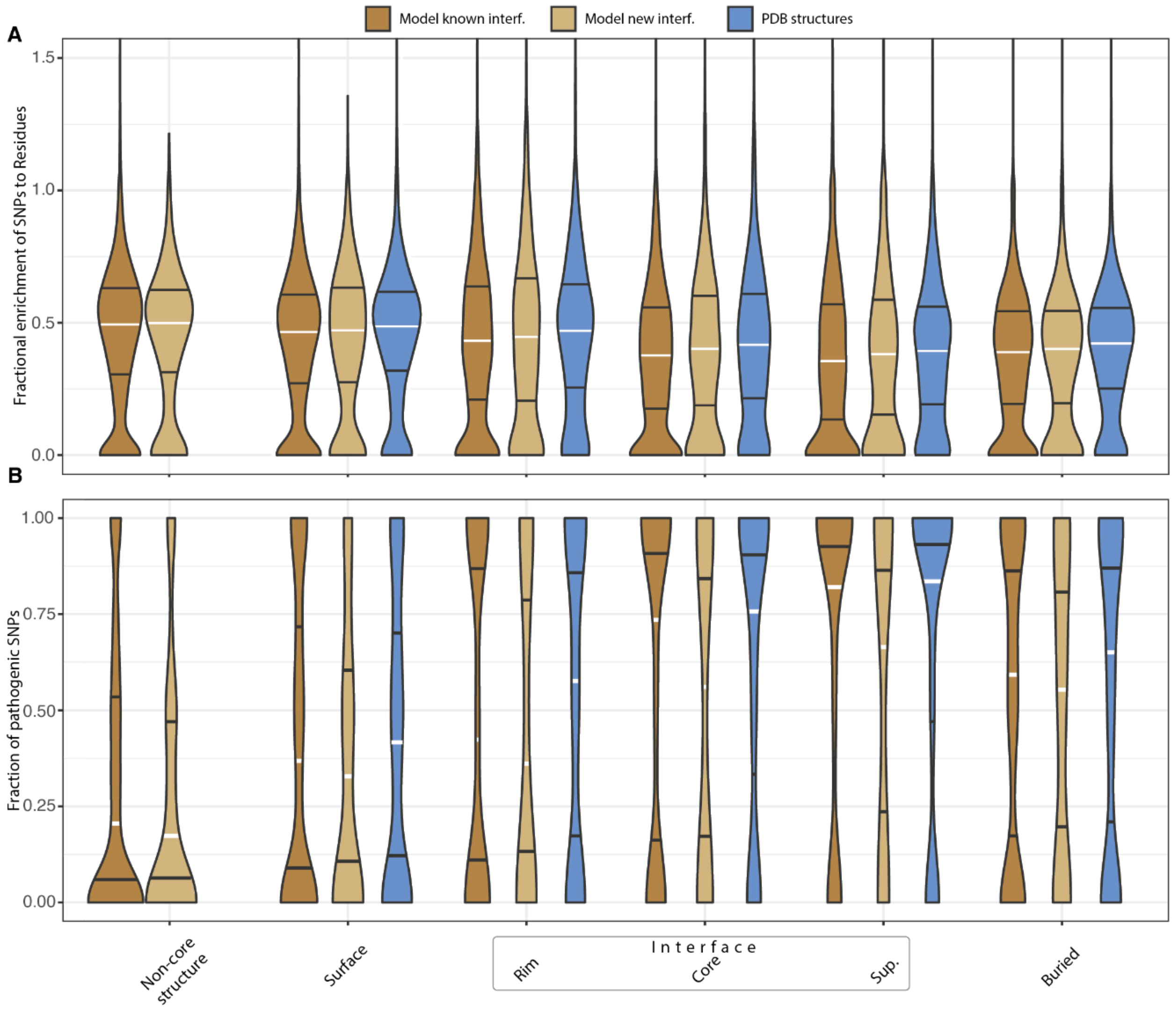


**Supplementary Figure 8. Distributions of SNPs and pathogenic SNPs in human proteins. A.** Violin plot depicting the fraction of SNPs [7] in each region relative to its size (number of residues). Three types of structures are compared: structures from the PDB (blue), models with quaternary structure types previously observed (dark brown) or novel (light brown). We calculated this ratio for each type and across protein regions as defined in [8]. An additional region, the "non-core structure", consists of residues absent from the core structure (Methods), which are enriched in residues with low pLDDT values. We note that one residue may contain up to 8 missense SNPs as different nucleotides of a codon can yield several types of missense mutations. Certain regions can contain more SNPs than residues, so the density extends beyond 1, although it approaches 0 at those values. We only show the range 0 to 1.5 for clarity. **B.** Violin plot showing the fraction of pathogenic SNPs in the same complex types and across the same regions. Clinical annotations are derived from ClinVar [9]. The number of pathogenic SNPs in each region is normalized by the number of benign and pathogenic SNPs (Methods). White and black horizontal bars show the median and 25/75$^{th}$ quantiles of each distribution, respectively.

**Supplementary Table 1. Effect size and statistical significance for differences of distributions from Fig. S5.** Top: comparison of distributions of SNP frequency shown in Fig. S5A. Bottom: comparison of pathogenic SNP frequencies shown in Fig. S5B.

**Comparing SNPs frequencies across structural regions**

| | Surface/Interf. core | Surface/Inter. rim | Interior/Interf. support |
|---|---|---|---|
| **Effect size** | | | |
| New QS type | 23.3% | 14.5% | 9.1% |
| Known QS type | 32.1% | 13.7% | 17.6% |
| Known structures | 22.8% | 6.3% | 13.2% |
| **P-value of the comparison** | | | |
| New QS type | <0.0001 | 0.0035 | 0.0068 |
| Known QS type | <0.0001 | <0.0001 | <0.0001 |
| Known structures | <0.0001 | 0.0329 | 0.0004 |

**Comparing pathogenic SNP frequency across structural regions**

| | Interf. core /surface | Interf. rim / surface | Interf. Support / Interior |
|---|---|---|---|
| **Effect size** | | | |
| New QS type | 71.6% | 36.7% | 25.0% |
| Known QS type | 64.7% | 29.1% | 35.6% |
| Known structures | 74.4% | 39.3% | 31.9% |
| **P-value of the comparison** | | | |
| New QS type | 0.0027 | 0.0535 | 0.109 |
| Known QS type | <0.0001 | 0.0001 | <0.0001 |
| Known structures | <0.0001 | <0.0001 | <0.0001 |
